# Proteomic profiling uncovers sexual dimorphism in the muscle response to wheel running exercise in the FLExDUX4 murine model of facioscapulohumeral muscular dystrophy

**DOI:** 10.1101/2025.03.15.639012

**Authors:** Yusuke Nishimura, Adam Bittel, Abhishek Jagan, Yi-Wen Chen, Jatin Burniston

## Abstract

FLExDUX4 is a murine experimental model of facioscapulohumeral muscular dystrophy (FSHD) characterized by chronic, low levels of leaky expression of the human full-length double homeobox 4 gene (DUX4-fl). FLExDUX4 mice exhibit mild pathologies and functional deficits similar to people affected by FSHD. Proteomic studies in FSHD could offer new insights into disease mechanisms underpinned by post-transcriptional processes. We used mass spectrometry-based proteomics to quantify the abundance of 1322 proteins in triceps brachii muscle, encompassing both male and female mice in control and free voluntary wheel running (VWR) in Wild-type (n=3) and FLExDUX4 (n=3) genotypes. We report the triceps brachii proteome of FLExDUX4 mice recapitulates key skeletal muscle clinical characteristics of human FSHD, including alterations to mitochondria, RNA metabolism, oxidative stress, and apoptosis. RNA-binding proteins exhibit a sex-specific difference in FLExDUX4 mice. Sexual dimorphism of mitochondrial protein adaptation to exercise was uncovered specifically in FLExDUX4 mice, where females increased, but males decreased mitochondrial proteins after a 6-week of VWR. Our results highlight the importance of identifying sex-specific diagnostic biomarkers to enable more reliable monitoring of FSHD therapeutic targets. Our data provides a resource for the FSHD research community to explore the burgeoning aspect of sexual dimorphism in FSHD.

**In Brief:** Nishimura et al. conducted proteomic analysis of triceps brachii muscle in the FLExDUX4 murine model of FSHD and verified FLExDUX4 mice recapitulate key skeletal muscle clinical characteristics of human FSHD, including disruptions to the mitochondrial proteome and proteins associated with RNA metabolism, oxidative stress, and apoptosis. RNA-binding proteins and the mitochondrial proteome response to exercise exhibited sexual dimorphism in FLExDUX4 mice. Specifically, females exhibited increases, whereas males exhibited decreases in mitochondrial protein abundance after 6-weeks voluntary wheel running.

**Highlights:**

- FLExDUX4 muscle proteome mirrors pathophysiology of FSHD patient myoblasts
- Mitochondrial proteins are more abundant in FLExDUX4 as compared to WT mice
- Severity of proteome disruption is greater in male than female mice
- RNA-binding proteins exhibit a sex-specific difference in FLExDUX4 mice
- Sex-specific mitochondrial proteome response to VWR in FLExDUX4 mice

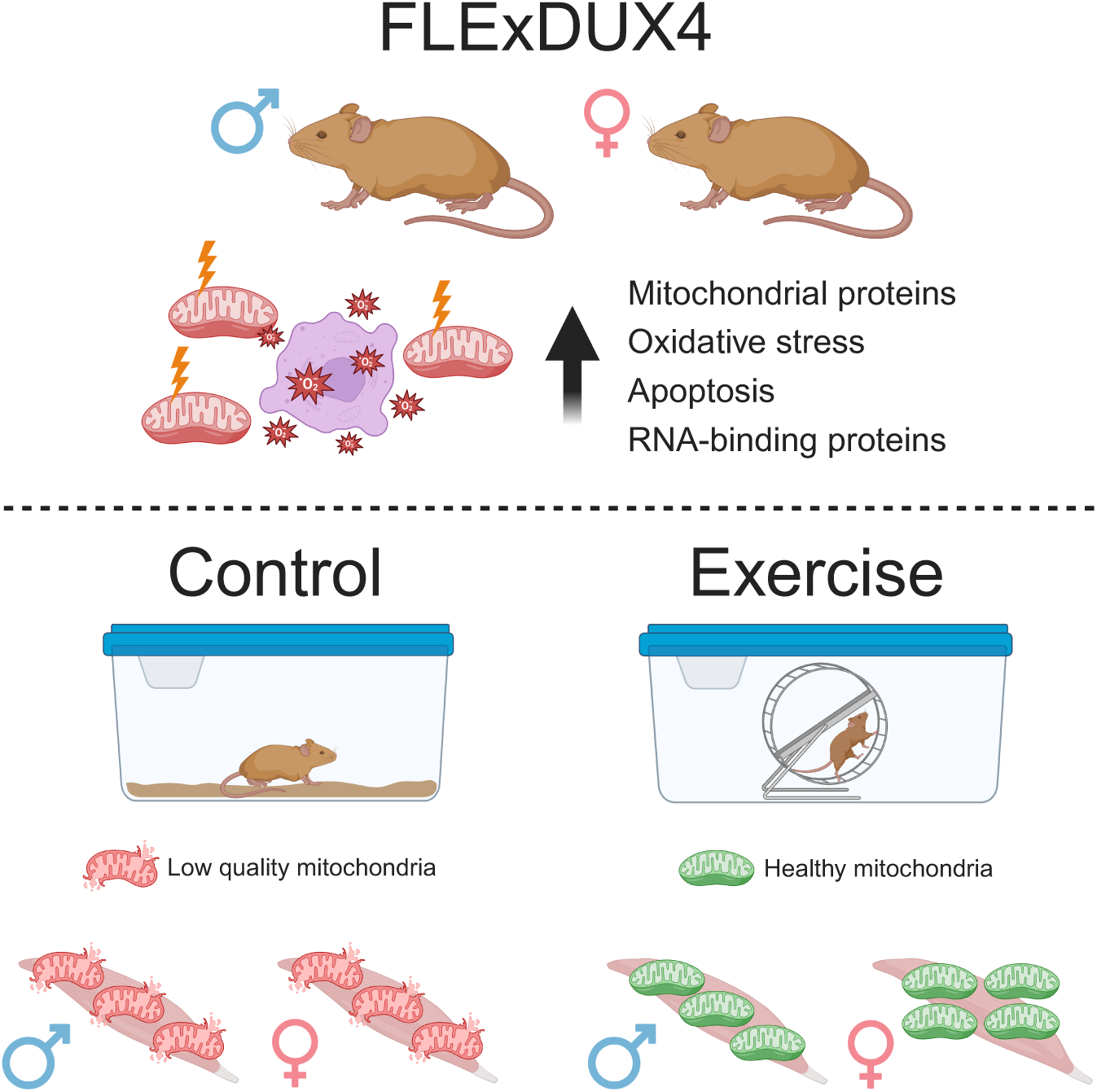

## Introduction

Facioscapulohumeral muscular dystrophy (FSHD) is a progressive autosomal dominant muscular dystrophy with an estimated prevalence of 1 in 8,000 to 1 in 20,000 worldwide and no known cure (1–3). FSHD is characterized by progressive and asymmetric weakness and wasting of muscles. Ectopic expression and activation of the double homeobox 4 (DUX4) gene is the primary molecular driver (4, 5) of FSHD and the disease is subcategorized into FSHD1 (OMIM 158900: ∼95 % of cases) and FSHD2 (OMIM 158901) depending on the genetic and epigenetic mechanisms of DUX4 activation. Clinical heterogeneity amongst people with FSHD is well established, including different ages of onset (6), and different rates of disease progression and severity (7). The clinical presentation of FSHD often occurs earlier in men than women (8), and magnetic resonance imaging reveals women with FSHD exhibit lesser muscle inflammation than men (9). The molecular basis for sexual dimorphism in FSDH is not yet understood and has been challenging to investigate due to the low and sporadic expression of DUX4 in skeletal muscles. Experimental systems including, i) muscle biopsies (10) or ii) myoblasts (11–13) from people affected by FSHD, iii) DUX4-induction via viral or vector infection in myoblasts (14, 15), and iv) preclinical animal models of FSHD (16) each have inherent challenges. DUX4 expression in patient samples is localised, varies on a temporal basis and likely occurs alongside secondary processes of inflammation and regeneration. Conversely, forced expression of DUX4 in experimental systems does not mimic the low and sporadic expression of DUX4 in patients’ muscles, and high-level DUX4 expression leads to rapid cell death, which complicates longer term studies. Despite the aforementioned challenges, decades of work have established DUX4-mediated downstream gene regulation (17, 18) and affected biological pathways in muscle, including DNA damage (11), p53-mediated apoptosis (19), disrupted RNA metabolism (20, 21), oxidative stress (22), impaired mitochondrial function (15) and structure (10), and losses in proteostasis (12, 13).

Jones et al. (23, 24) reports transgenic mice carrying full-length human DUX4 transgene (DUX4-fl) inverted and flanked by lox sites to bypass the embryonic lethality of DUX4 expression. Low expression of DUX4 mRNA occurs in adult FLExDUX4 transgenic animals (23) and, when crossed with muscle-specific Cre (ACTA1-MCM) mouse, ACTA1-MCM/FLExDUX4 bi-transgenic animals exhibit muscle-specific expression of DUX4 and mild FSHD-related pathology (24). Moderate and severe models of FSHD pathology can be achieved by graded tamoxifen treatment in ACTA1-MCM/FLExDUX4 bi-transgenic animals and are associated with decrements in exercise capacity and muscle function (24). After moderate tamoxifen treatment, the exercise capacity of ACTA1-MCM/FLExDUX4 animals decays and is restored within a period of ∼30 days, whereas higher levels of tamoxifen treatment result in more severe muscle dysfunction and it is necessary to euthanise animals less than 10 days after tamoxifen treatment (24). Such abrupt and transient responses to DUX4 activation provide useful bioassays for rapidly assessing DUX4-targetted therapeutic interventions but lack application to the chronic, low-level expression of DUX4 that people affected by FSHD live with throughout adulthood.

Murphy et al. (25) characterises the FLExDUX4 single transgene mice that exhibit low-level ‘leaky’ DUX4 expression and, across a period of 12 months, experience changes in muscle fibre profile and losses in muscle mass and function. Male FLExDUX4 mice exhibit muscle dysfunction at a younger age than in females (25), which recapitulates the earlier clinical presentation of FSHD in men than women (3). Furthermore, despite similar levels of DUX4 gene expression between sexes, male FLExDUX4 mice exhibit greater muscle fibrosis than females (26). A 6-week period of voluntary wheel running (VWR) improves the molecular and functional deficits in male FLExDUX4 mice (26) but the molecular responses to exercise in females were not investigated. Based on bioinformatic analyses of RNAseq data, signalling pathways involved in actin cytoskeletal, inflammation, fibrosis, and muscle wasting are altered in male FLExDUX4 animals (26). VWR ameliorated transcriptional changes of the impaired signalling pathways, but it is unknown whether these changes at the transcript level are translated to the muscle proteome. In human myoblasts, DUX4 expression disrupts the expected connection between mRNA and protein responses (14) and in myoblasts from FSHD patients there is a disconnection between the mRNA expression (15) and protein abundance (13) of mitochondrial proteins that may be due to an accumulation of mitochondrial proteins with lower than expected turnover rates (13).

Proteomic studies in FSHD are currently underrepresented so further exploitation of proteomics offers access to an, as yet, untapped resource of new Information. Herein, we report proteomic analysis of samples from (26) and use multi-factor experimental designs to investigate the interactions between genotype (FLExDUX4 vs Wild-type) sex (male vs female) and training status (VWR vs sedentary). Consistent with our findings in myoblasts from FSHD patients (13), mitochondrial proteins were more abundant in FLExDUX4 compared to WT mice. In response to VWR, male FLExDUX4 mice decreased, whereas female FLExDUX4 mice increased mitochondrial protein abundance, indicating sexual dimorphism of mitochondrial proteome adaptation to exercise. Together, these findings highlight the importance of investigating sex-specific diagnostic biomarkers and therapeutic interventions in FSHD.

## Experimental Procedures

### FLExDUX4 animal models

**Figure 1 A** provides an overview of the experimental design and protocol, which consists of a 3-factor design to investigate interactions between genotype (WT versus FLExDUX4), sex (Male versus Female), and training status (control versus exercise) using n = 3 mice (n = 24, total). We have previously characterized the physiological phenotype of male and female FLExDUX4 mice and their C57BL/6 wild type (WT) littermates (26). A subset of mice with physiological characteristics consistent (**Supplemental Fig. 1**) to those previously reported (26) was used for the current proteomic analysis. All mice were handled in accordance with the guidelines established by the Institutional Animal Care and Use Committee (IACUC) of the CNMC in Washington, D.C., and all procedures were carried out under the approved animal protocol. A total of 24 triceps brachii muscles was used for proteomic analyses. Male and female FLExDUX4 mice and their C57BL/6 wild type (WT) littermates were randomly assigned to voluntary, non-resisted wheel running (Exercise), or a no-wheel-running control condition (Control). The triceps brachii was selected because it is affected early and consistently in people with FSHD (27) and triceps brachii demonstrates a high muscle activity profile during running in rodents (28). Mice in the training group were provided 24-hour access to an 11 cm wheel for 40 days, while the control group remained in cages without a wheel. The FLExDUX4 mouse model of FSHD expresses a human DUX4-fl transgene in a C57BL/6 background under the control of the Rosa26 promoter after Cre-recombinase-mediated inversion (23). In the absence of Cre-recombinase, these mice present with low levels of “leaky” DUX4 mRNA expression through antisense transcription consistent with the levels of DUX4 mRNA measured in human FSHD myocytes (23). We previously reported similar evidence for leaky DUX4 mRNA in our FLExDUX4 male and female mice (26).

**Figure 1.**
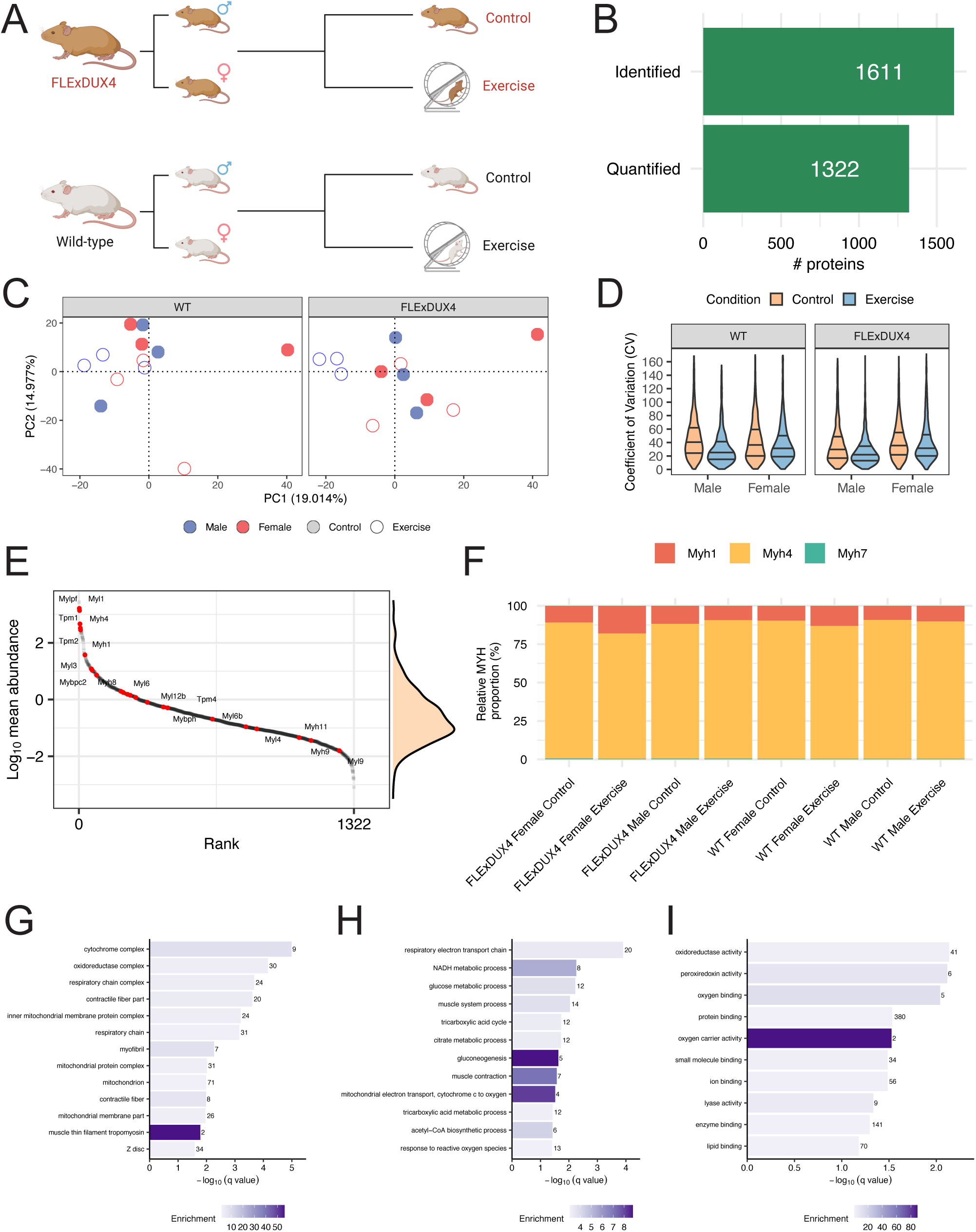
Proteomic profiling to uncover sexual dimorphism and exercise responses of triceps muscle of FLExDUX4 mice. (***A***) Overview of the experimental design. A 3-factor design to investigate interactions between Genotype (WT versus FLExDUX4), Sex (Male versus Female), and training status (Control versus Exercise) using n = 3 mice (n = 24). (***B***) The number of proteins confidently identified (1% < FDR) at least n = 1 and quantified in all n = 24 samples. (***C***) Principal Component Analysis (PCA) on protein abundance. (***D***) Violin plot comparing the coefficient of variation in protein abundance between n = 3 mice within an experimental group was calculated on a protein-by-protein basis. (***E***) Rank distribution plot of average protein abundance quantified (n = 1322 proteins). Myosin-associated proteins are highlighted in red data point and annotated. (***F***) Stacked bar chart illustrating a relative proportion of Myh1 (red), Myh4 (yellow), and Myh7 (green) to compare between experimental groups. Gene ontology terms in (***G***) Cellular Component, (***H***) Biological Process, and (***I***) Molecular Function enriched in a whole proteome quantified in this study (n = 1322 proteins).

### Protein extraction and quantification

Proteins were extracted from muscle samples as previously described (29, 30). Muscle samples were ground in liquid nitrogen using a Cellcrusher (Cellcrusher, Cork, Ireland), then homogenized on ice in 10 volumes of 1 % Triton X-100 (v/v), 5 mM NEM, 5 mM EDTA, 50 mM Tris, pH 7.4 including phosphatase inhibitor and complete protease inhibitor cocktails (Roche Diagnostics, Lewes, United Kingdom) using a PolyTron homogenizer (KINEMATICA, PT 1200 E) followed by sonication (Fisherbrand^TM^). Homogenates were incubated on ice for 15 min, then centrifuged at 1000 x *g*, 4 °C, for 5 min to fractionate myofibrillar (pellet) from soluble (supernatant) proteins. Myofibrillar proteins were resuspended in a half-volume of homogenization buffer followed by centrifuged at 1000 x g, 4 °C, for 5 min. The washed myofibrillar pellet was then solubilized in lysis buffer (24 mM sodium lauroyl sarcosinate, 40 mM sodium deoxycholate, 100 mM Tris-HCl pH 9.0 including phosphatase inhibitor and complete protease inhibitor cocktails). Aliquots of protein were precipitated in 5 volumes of ice-cold acetone and incubated overnight at -20 °C then pelleted and resuspended in 300 μL of lysis buffer. TCEP (Bond Breaker™) was added to each sample (5 mM final concentration) and incubated at 95 °C for 30 min using a water bath. Samples were cooled to RT and chloroacetamide was added to each sample (20 mM final concentration) and incubated at 4 °C for 30 min. Samples were centrifuged at 12,000 x g, 4 °C for 20 min and supernatant was stored as myofibrillar fraction. Total protein concentration (μg/μl) was quantified against bovine serum albumin (BSA) standards using the Bradford assay (Thermo Scientific, #23236), according to the manufacturer’s instructions.

### Protein digestion

Tryptic digestion was performed using the filter-aided sample preparation (FASP) method (31). Aliquots containing 50 µg protein were washed with 200 µl of UA buffer (8 M urea, 100 mM Tris, pH 8.5). Proteins were incubated at 37 °C for 15 min in UA buffer containing 100 mM dithiothreitol followed by incubation (20 min at 4 °C) protected from light in UA buffer containing 50 mM iodoacetamide. UA buffer was exchanged for 50 mM ammonium bicarbonate and sequencing-grade trypsin (Promega, V5113, Madison, WI, USA) was added at an enzyme to protein ratio of 1:50. Digestion was allowed to proceed at 37 °C overnight then peptides were collected in 100 μl 50 mM ammonium bicarbonate containing 0.2 % (v/v) trifluoroacetic acid. Samples containing 5 µg of peptides were de-salted and eluted in 40 % ACN, 0.1 % TFA via Oasis HLB 96-well µElution Plate, 2 mg Sorbent per Well, 30 µm Particle Size (Waters, 186001828BA), and the peptides were dried by vacuum centrifugation (Thermo Scientific, SpeedVac SPD1030). Peptides were resuspended in 25 µl of loading buffer (2 % (v/v) acetonitrile, 0.1 % (v/v) formic acid) containing 10 fmol/ μl yeast ADH1 (MassPrep standard, 186002328, Waters Corp., Milford, MA) in preparation for LC-MS/MS analysis.

### Liquid chromatography-mass spectrometry analysis

Data-dependent analysis of digests from muscle protein fractions was performed using an Ultimate 3000 RSLCTM nano system (Thermo Scientific) coupled to a Q-Exactive orbitrap mass spectrometer (Thermo Scientific). Samples (2.5 μL corresponding to 500 ng of protein) were loaded on to the trapping column (Thermo Scientific, PepMap NEO, 5 μm C18, 300 μm X 5 mm), using µlPickUp injection, for 1.5 minute at a flow rate of 25 μl/min with 0.1 % (v/v) TFA and 2 % (v/v) ACN. Samples were resolved on a 110 cm micro pillar array column (uPAC™ Neo; 180 μm bed width, 16 μm pillar length, 2.5 μm pillar diameter and 300 Å pore size grafted with C_18_ groups) using a 105-minute gradient, starting with 97.5 % A (0.1% formic acid in water) and 2.5 % B (79.9 % ACN, 20 % water, 0.1 % formic acid) at a flow rate of 0.75 µL/min, increasing to 3 % B at 1.0 min, 9.4 % B at 1.1 min, 14.6 % B at 8.9 min, and 14.8 % B at 9.0 min, then 28 % B at 69.0 min, 56 % B at 105.0 min, and finally 95 % B at 105.5 min, with the flow rate reducing to 0.25 µL/min from 9.0 min onward. Following the gradient, the column was washed with 95 % B at 108.5 min for 0.5 minutes, followed by a re-equilibration to 5 % B at 109 min and holding until 112 min. Another wash cycle with 95 % B was performed at 112.5 min, held until 115.5 min, followed by re-equilibration at 5 % B from 116 min to 119 min. The final wash with 95 % B occurred at 119.5 min until 122.5 min, followed by a re-equilibration to 3 % B at 123 min, held until 130 min, before increasing the flow rate to 0.75 µL/min at 131 min, and holding until 145 min to ensure the column is fully equilibrated and ready for the next injection. The data-dependent selection of the top-10 precursors selected from a mass range of m/z 300-1600 consisted of a 70,000-resolution (at m/z 200) MS scan (AGC set to 3^e6^ ions with a maximum fill time of 240 ms). MS/MS data were acquired using quadrupole ion selection with a 3.0 m/z window, HCD fragmentation with a normalized collision energy of 30 and orbitrap analyzer at 17,500 resolution at m/z 200 (AGC target 5^e4^ ion with a maximum fill time of 80 ms). To avoid repeated selection of peptides for MS/MS, the program used a 30 s dynamic exclusion window.

### Label-free quantitation of protein abundances

Progenesis Quantitative Informatics for Proteomics (QI-P; Nonlinear Dynamics, Waters Corp., Newcastle, UK, Version 4.2) was used for label-free quantitation consistent with our previous work (13, 30, 32). Prominent ion features were used as vectors to warp each data set to a common reference chromatogram. An analysis window of 8–112 min and 300–1600 m/z was selected. Log-transformed MS data were normalized by inter-sample abundance ratio, and relative protein abundances were calculated using unique peptides only. Abundance data were then normalised to the 3 most abundant peptides of yeast ADH1 to derive abundance measurements in fmol/ μg protein (33). MS/MS spectra were exported in Mascot generic format and searched against the Swiss-Prot database restricted to Mus musculus (house mouse) (17,089 sequences) using a locally implemented Mascot server (v.2.8; www.matrixscience.com). The enzyme specificity was trypsin with 2 allowed missed cleavages, carbamidomethylation of cysteine (fixed modification) and oxidation of methionine (variable modification). m/z error tolerances of 10 ppm for peptide ions and 20 ppm for fragment ion spectra were used. Peptide results were filtered to 1% FDR based on decoy search and at least one unique peptide was required to identify each protein. The Mascot output (xml format), restricted to non-homologous protein identifications was recombined with MS profile data in Progenesis.

### Experimental design and statistical rationale

This study investigates differences in the muscle proteome between male and female FLExDUX4 and wild-type mice under control conditions or in response to 6 weeks VWR exercise. Our experiment encompasses 3 independent factors of (i) genotype (WT versus FLExDUX4), (ii) sex (Male versus Female), and (iii) training status (control versus exercise). Interactions across 3-factor designs can be challenging to interpret, particularly in the context of omic data, therefore, subsets of experimental questions were investigated using separate 2-factor analyses of variance (ANOVA). Our primary aim was to investigate sex-specific and common effects of the FLExDUX4 transgene, which was tested by two-way ANOVA on Genotype (FLExDUX4 vs WT) and Sex (Male vs Female), in animals that did not perform VWR. Our secondary aim was to investigate sex-specific responses to VWR in FLExDUX4 mice, and this was achieved through separate 2-factor ANOVA comparing training status (Control vs. VWR) and Sex (Male vs Female) in either FLExDUX4 or WT mice. All statistical analysis was performed in R (Version 4.3.2). Differences in the abundance of proteins between FLExDUX4 and WT or Exercise and Control groups are reported as log_2_ transformed data and statistical significance was set at P < 0.05. Due to the limited number of replicates (n = 3, per group) a false discovery rate criterion was not set, instead, q values (34) at the p = 0.05 threshold were reported.

### Bioinformatic analysis

Gene ontology analysis (GO) was performed via Overrepresentation Enrichment Analysis (35) using Gene Ontology enRIchment anaLysis and visuaLizAtion tool (Gorilla) (36). Enrichment of GO terms was considered significant if the Benjamin Hochberg adjusted p value was 0.01. The coverage of mitochondrial proteins was surveyed against Mouse MitoCarta 3.0 (37).

## Results

### Proteomic profiling of triceps brachii muscle

Proteomic analysis was conducted on 24 samples of triceps brachii muscle, encompassing both male and female WT and FLExDUX4 mice in control and exercised conditions (**Fig. 1 A**). Label free quantitation identified 1611 proteins (>1 unique peptide and <1% FDR) across soluble and myofibrillar fractions (**Supplemental Fig. 2 A**). After filtering to exclude missing values (**Supplemental Fig. 2 B**), 1322 proteins were quantified in all n=24 samples (**Fig. 1 B**). Missing values in proteomics experiments may represent biologically relevant information but we did not find patterns that indicated clear biological differences between groups, therefore, we interpret missing data in this experiment as being primarily due to inconsistent detection of low abundance proteins. Consistent with our earlier proteomic study on myoblasts from people affected by FSHD (13), we did not detect previously suggested DUX4 target candidate genes highlighted by Yao et al. (17). Principal component analysis on muscle proteome data separated the Control and Exercise groups in WT females and FLExDUX4 males (**Fig. 1 C**). The median and inter-quartile range in coefficient of variation (CV) amongst protein abundances in each group of n = 3 mice was M = 30.4 % (Q1 17.9 % to Q3 50.3%, **Fig.1 D**). Protein abundances spanned from minimum of log_10_ -3.09 (fmol/μg of protein) to maximum of log_10_ 3.44 (fmol/μg of protein) (**Fig. 1 E**). Fast-isoforms of myosin heavy chains, Myh1 (MyHC-2x) and Myh4 (MyHC-2b), were more abundant than the slow Myh7 isoform (MyHC-slow) (**Fig. 1 E & F**) but the relative proportion of Myh isoforms did not differ (Three-way ANOVA P > 0.05) across groups. Bittel et al. (26) reports the proportion of Myn2 increased in exercised WT and FLExDUX4 males but Myh2 (MyHC-2a) was not distinguished from Myh4 in our proteomic data. In mouse Myh2 and Myh4 share 96 % similarity and no tryptic peptides were specific to Myh2 were detected. Gene ontology (GO) analysis of Cellular Component (**Fig. 1 G**), Biological Process (**Fig. 1 H**), and Molecular Function (**Fig. 1 I**) of the proteome quantified in this study (n = 1332 proteins) was significantly enriched with contractile fiber part (GO:0044449), myofibril (GO:0030016), mitochondrion (GO:0005739) and glycolytic process (GO:0006096), which are typical of fast-twitch skeletal muscle.

### Proteomic features of FLExDUX4 mice recapitulate key skeletal muscle clinical characteristics of people affected by FSHD and sex-specific differences

Control mice that did not perform VWR exhibited 9 statistically significant interactions between genotype (FLExDUX4 vs WT) and sex (Male versus Female) and a further 55 proteins exhibited statistically significant differences between genotype that were common to both male and female animals (**Fig. 2, Supplemental Table S1**). The majority (50 of 55 proteins) exhibiting a main effect of genotype were more abundant in FLExDUX4 compared to WT (purple data points in the top right quadrant, **Fig. 2 A**) and were enriched in gene ontology terms, including Mitochondrion and Mitochondrial protein complex. In all, our data included 268 of the 1140 proteins annotated in the Mouse MitoCarta database (**Supplemental Fig.3 A**), and 21 mitochondrial proteins were significantly more abundant in FLExDUX4 compared to WT mice (P < 0.05 main effect of genotype, **Fig. 2 B**), including proteins of the electron transport chain, oxidative phosphorylation, mitochondrial ribosome, mitochondrial metabolism and mitochondrial dynamics. Manual data interrogation found other common features, including indicators of mitochondrial dysfunction, oxidative stress, apoptosis, and alterations in iron storage (**Fig. 2 B, Supplemental Fig. 4**) amongst the proteins that exhibited significant differences between WT and FLExDUX4 muscles.

**Figure 2.**
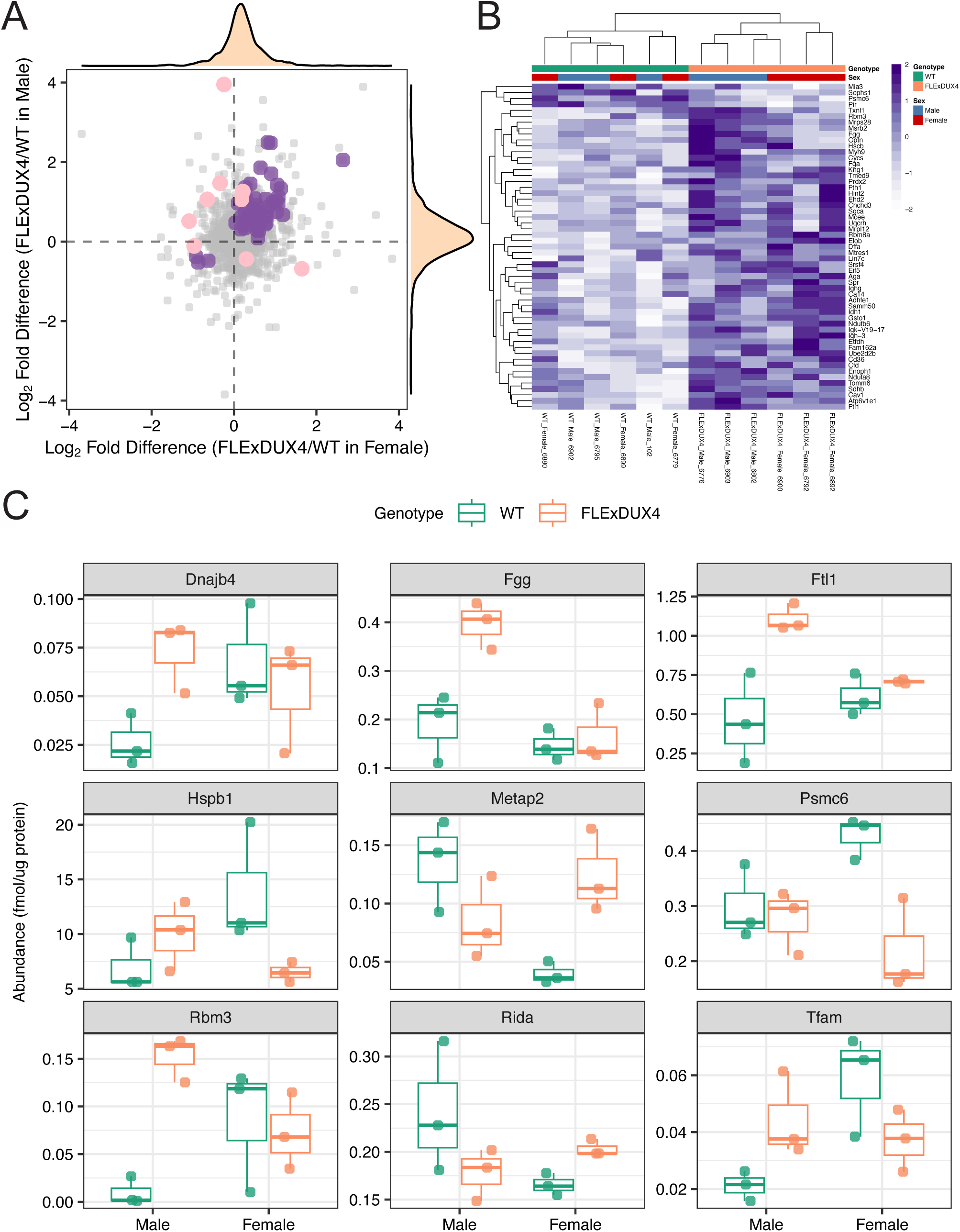
Proteomic profiling uncovers sexual dimorphism of FLExDUX4. (***A***) Scatter plot comparing the differences in the Log_2_ Fold-Difference (FLExDUX4/WT) between protein abundance of females (x-axis) and males (y-axis) in control mice. Purple data points represent proteins with a main effect of Genotype (P < 0.05) and pink data points represent proteins with an interaction effect between Genotype and Sex (P < 0.05). Density plots display the distribution of individual protein abundance data both in X and Y axis. (***B***) Heatmap of proteins with significant differences in abundance between FLExDUX4 and WT muscle (proteins, a main effect of Genotype; P < 0.05). Protein abundances were normalized by row and represented by colors according to the scale in the key. Both rows (proteins) and columns (samples) were clustered using hierarchical clustering. (***C***) Boxplot of 9 proteins that exhibit an interaction effect (P < 0.05, two-way ANOVA of genotype and sex in control mice).

The 9 proteins exhibiting significant interactions between genotype and biological sex (**Fig. 2 C)** provide new insight into sexual dimorphism in the muscle response to DUX4 expression. Methionine aminopeptidase 2 (Metap2) is an inhibitor of kinases that phosphorylate eukaryotic initiation factor 2 alpha subunit (eIF2-α) (38) and was less abundant in Male FLExDUX4, but more abundant in Female FLExDUX4 compared to their respective WT controls (**Fig. 2 C**). Phosphorylation of eIF2-α is associated with inhibition of translation during cellular stress responses, including the unfolded protein response, and our data at the protein level adds to findings (39) reporting Metap2 gene expression is elevated in FSHD myoblasts and differentiated FSHD myotubes. RNA-binding protein 3 (Rbm3) is a cold-induced member of the glycine-rich RNA-binding protein family and was specifically more abundant in male FLExDUX4 mice (**Fig. 2 C**). Several RNA-binding proteins, including Rbm3 were previously identified as DUX4/4c binding partners (40), and Rbm3 contains an RNA-recognition motif and, therefore, may have a regulatory or selective role in translation. RNA-binding protein 8A (Rbm8a) is a component of the spliceosome and required for pre-mRNA splicing. Jagannathan et al. (14) reported that induction of DUX4 in immortalised MB135 human myoblasts reduced Rbm8a at the mRNA level, whereas the protein abundance of Rbm8a was unchanged. In our proteomic data, Rbm8a was specifically more abundant in FLExDUX4 mice (P=0.0147), but an interaction effect was not evident (P=0.19) (**Fig. 2 B**). DnaJ homolog subfamily B member 4 (Dnajb4) was more abundant in FLExDUX4 males compared to WT while there were no differences between females (**Fig. 2 C**). DnaJ homolog subfamily B member 4 (Dnajb4) is a cochaperone of the ubiquitous Hsp70 chaperone and the loss of Dnajb4 function is known to cause a myopathy with early respiratory failure due to a possible aggregation of Dnajb4 client proteins (41). Transcription Factor A, mitochondrial (Tfam) is a nuclear-encoded protein that plays a role in controlling mtDNA copy number (42). Tfam was more abundant in FLExDUX4 mice than WT in male, but Tfam was less abundant in FLExDUX4 mice than WT in female. Similarly, heat shock protein beta-1 (Hspb1) was more abundant in FLExDUX4 mice than WT in male, but Hspb1was less abundant in FLExDUX4 mice than WT in female. Collectively, these proteins represent cellular stress response mechanisms, including the regulation of mRNA stability, protein folding, and the cellular response to heat shock and oxidative stress, which may point to a sex-specific difference in FLExDUX4 mice.

### Sexual dimorphism of mitochondrial proteome adaptation to VWR in FLExDUX4 mice

In FLExDUX4 mice, significant interactions (P<0.05, two-way ANOVA) between training status (Control versus VWR) and sex (Male versus Female) were evident for 48 proteins (**Supplemental Table S2**). The majority (29 of 48 proteins) increased in females but decreased in males exposed to VWR (pink data points in the bottom right quadrant, **Fig. 3 A**) and gene ontology terms, including mitochondrion and mitochondrial respiratory chain complex were enriched amongst these 29 proteins (bottom right quadrant; **Fig. 3 C**). In contrast, 14 proteins decreased in females, but increased in males (pink data points in the top left quadrant, **Fig. 3 A**). No gene ontology terms were significantly enriched but Metap2 was amongst the 14 proteins in the top left quadrant and was more abundant in FLExDUX4 female but less abundant in male FLExDUX4 as compared to their respective WT control (**Fig. 2 B, Supplemental Fig. 4**). Three-way ANOVA of genotype, sex, and training status on the average mitochondrial protein abundance (268 proteins) did not find significant differences, but exercise tended to increase the average mitochondrial protein abundance in female FLExDUX4 mice while the mitochondrial protein abundance in male FLExDUX4 mice remained unchanged (**Supplemental Fig.3 B**). Ninety-six proteins exhibited differences between VWR and control that were common to both male and female FLExDUX4 mice (P<0.05, main effect of training status). Proteins associated with oxidative stress and redox homeostasis, including Gsr, Gstp1, Gstm1, Selenbp1, Aldh4a1, Aldh6a1, Aldh7a1 were decreased by exercise similarly in both male and female FLExDUX4 mice (**Fig. 3 B, Supplemental Fig. 5**) and proteins associated with proteostasis, including Hspb8, Pdia6 were commonly increased by exercise. In addition, Samm50, which was significantly more abundant in FLExDUX4 compared to WT control in the non-exercised condition, was decreased by exercise in FLExDUX4 mice (**Fig. 2 B, Supplemental Fig. 4**).

**Figure 3.**
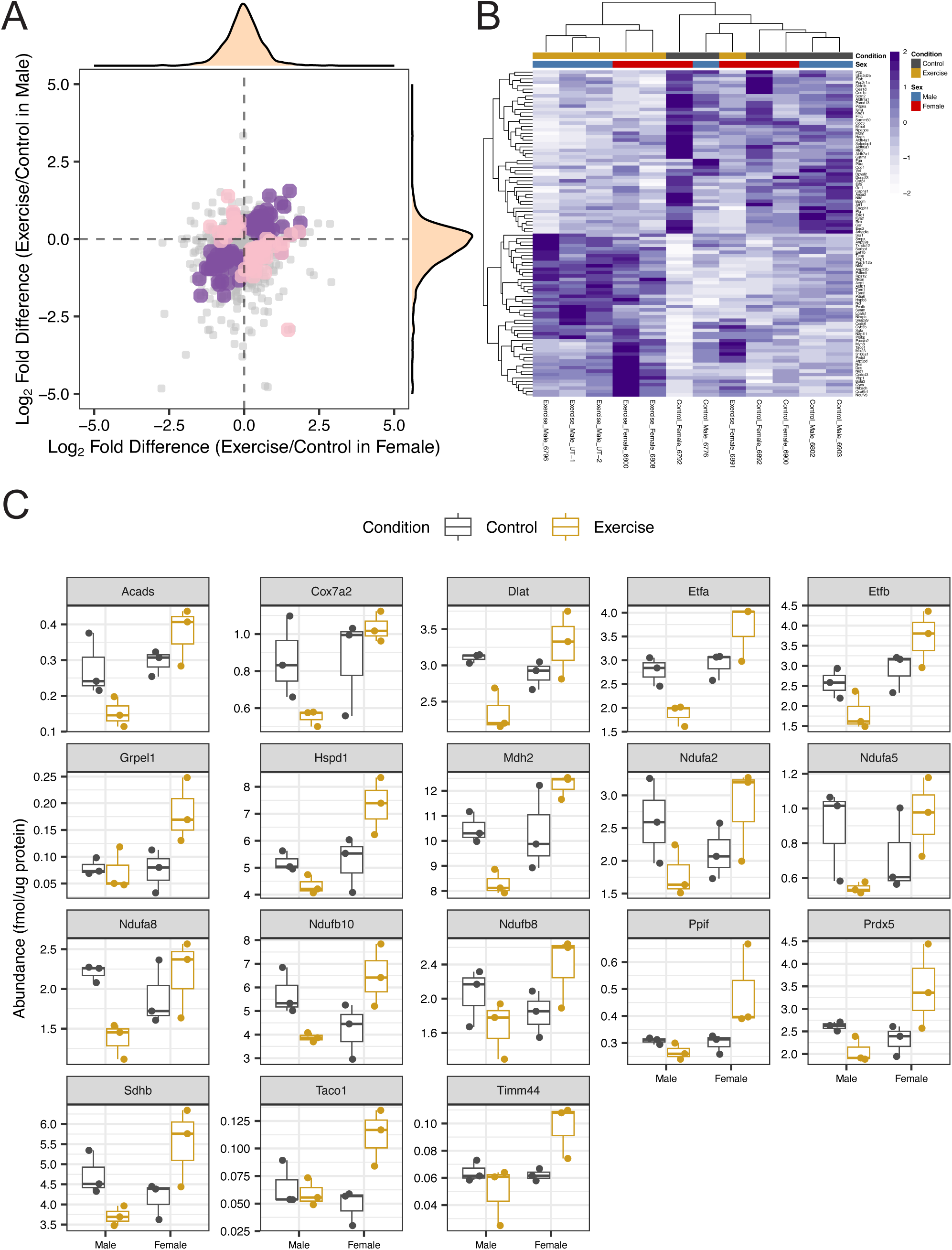
Altered mitochondrial proteome responses to VWR between males and females in FLExDUX4. *(**A**)* Scatter plot comparing the differences in the Log_2_ Fold-Difference (Exercise/Control) between protein abundance of females (x-axis) and males (y-axis) in FLExDUX4 mice. Purple data points represent proteins with a main effect of Exercise (P < 0.05) and pink data points represent proteins with an interaction effect between training status and Sex (P < 0.05). Density plots display the distribution of individual protein abundance data both in X and Y axis. (***B***) Heatmap of proteins with significant differences in abundance between Exercise and Control muscle (proteins, a main effect of training status; P < 0.05). Protein abundances were normalized by row and represented by colors according to the scale in the key. Both rows (proteins) and columns (samples) were clustered using hierarchical clustering. (***C***) Boxplot of 18 mitochondrial proteins filtered by Mouse MitoCarta that are localized in the bottom right quadrant with an interaction effect (P < 0.05, two-way ANOVA of training status and Sex in FLExDUX4 mice).

### Mitochondrial proteome responses to VWR in wild-type skeletal muscle

Interactions between training status (Control versus Exercise) and biological sex were also evident in WT mice (**Supplemental Fig. 7 A & Supplemental Fig. 8**) but the proteins that differed by sex amongst WT animals had little overlap with the aforementioned proteins that exhibited sex-specific responses amongst FLExDUX4 animals (**Supplemental Table S3**). Only 4 (Atp5pd, Hspb8, Pdlim5, Pzp) of the 97 proteins that exhibited significant main effect of training status in WT animals were also highlighted in equivalent analysis of FLExDUX4 mice. In WT mice, VWR was associated with a greater abundance of mitochondrial proteins in both male and female WT mice, which contrasts with the sex-specific responses in FLExDUX4 animals. Mitochondrial proteins were enriched amongst the 68 proteins (out of 97 proteins, **Supplemental Fig. 7 B**) that exhibited a main effect of training status and were increased in response to exercise both in male and female WT mice (**Supplemental Fig. 7 B**). Five mitochondrial proteins (Ak3, Fxn, Gcsh, Mmab, Msrb3) exhibited sex-specific responses in WT but these were discrete from the proteins that exhibited interaction effects in FLExDUX4 mice (**Supplemental Fig. 7 C**).

### VWR reversed altered proteomic features of FLExDUX4 mice in both males and females

In male mice 37 proteins exhibited statistically significant interactions between genotype (WT versus FLExDUX4) and training status (Control versus Exercise), including 27 proteins that were more abundant in FLExDUX4 as compared to WT in the non-exercised condition and were decreased to the WT control levels by VWR in FLExDUX4 (pink data points in the bottom right quadrant, **Fig. 4 A & B, Supplemental Table S4**). Ten out of those 27 proteins were found in Mouse MitoCarta, including proteins involved in mitochondrial respiratory chain complex. Proteins involved in proteostasis (Hspa1a, Btf3) and iron storage (Ftl1, Tf) were also found in the bottom right quadrant. Btf3 associates with the nascent polypeptide-associated complex, and prevents inappropriate targeting of non-secretory polypeptides to the endoplasmic reticulum (ER). Hnrnph1 was more abundant in FLExDUX4 and VWR further increased the abundance (**Supplemental Fig. 9**). Hnrnph1 is a component of the heterogeneous nuclear ribonucleoprotein (hnRNP) complexes which provide the substrate for the processing events that pre-mRNAs undergo before becoming functional, translatable mRNAs in the cytoplasm.

**Figure 4.**
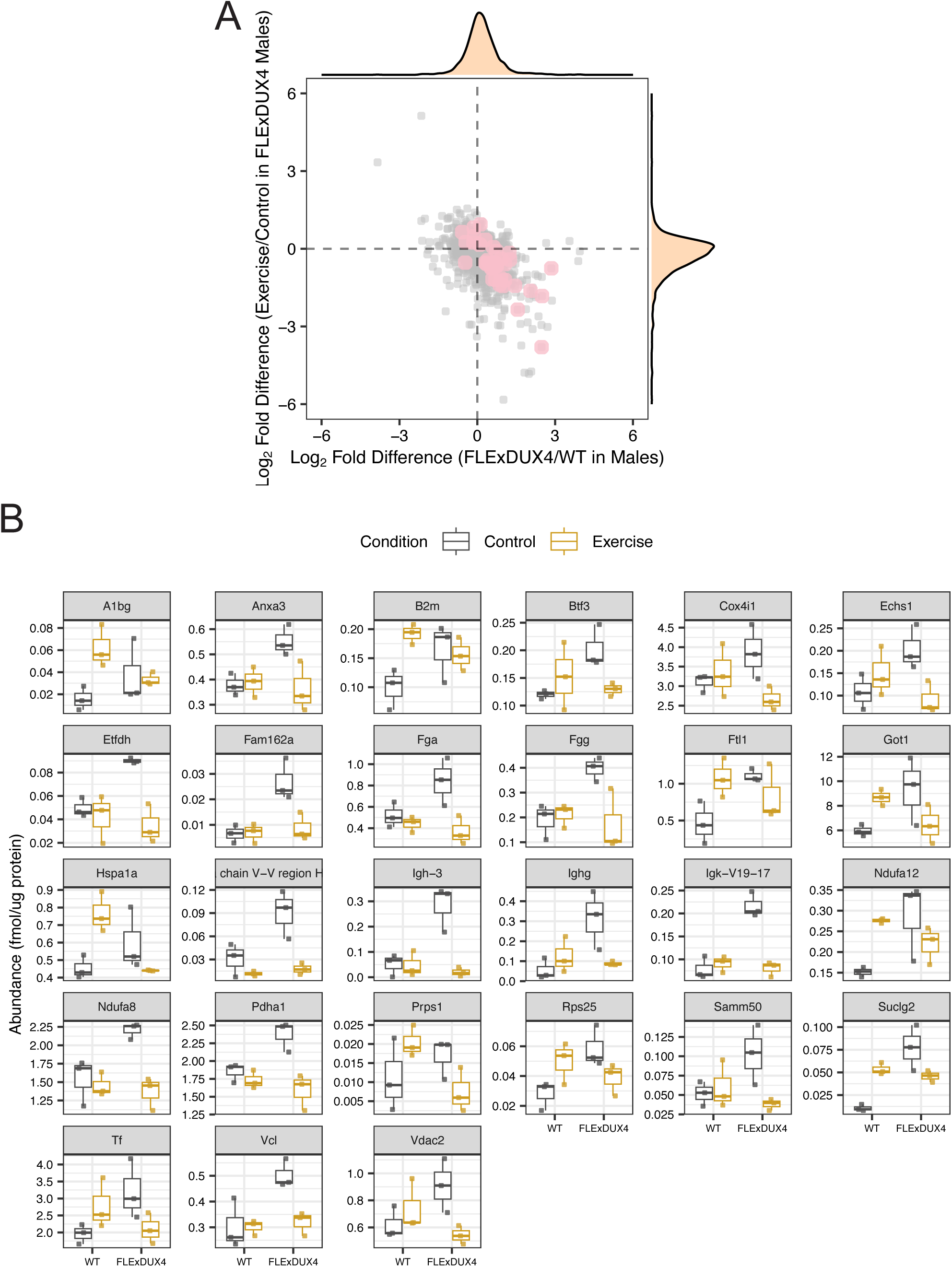
VWR normalized altered mitochondrial protein abundance in male FLExDUX4 mice. *(**A**)* Scatter plot comparing the Log_2_ Fold-Difference between FLExDUX4 control and WT control (x-axis) and the Log_2_ Fold-Difference between FLExDUX4 Exercise and FLExDUX4 control (y-axis). Pink data points represent proteins with an interaction effect between training status and Genotype (P < 0.05) in male mice. Density plots display the distribution of individual protein abundance data both in X and Y axis. (***B***) Boxplot of 27 proteins that are localized in the bottom right quadrant with an interaction effect (P < 0.05, two-way ANOVA of training status and Genotype in male mice).

Two-way ANOVA of genotype (WT versus FLExDUX4) and training status (Control versus Exercise) in female mice exhibits 27 proteins with statistically significant interaction (**Fig. 5, Supplemental Table S5**). Among 27 proteins with a significant interaction effect (**Supplemental Fig. 10**), 21 proteins were more abundant in FLExDUX4 as compared to WT, but decreased to the WT control levels by VWR (pink data points in the bottom right quadrant, **Fig. 5 A & B**). Four (Spr, Rida, Mmab, Gsr) out of those 21 proteins were found in Mouse MitoCarta, but are involved in iron storage (Ftl1) and the ubiquitin proteasome system (Otud6b, Uchl3, Sugt1, Elob) rather than mitochondrial respiration. In contrast, Txndc12 was less abundant in FLExDUX4 as compared to WT but increased by VWR (**Supplemental Fig. 10**). Txndc12 is protein-disulfide reductase of the endoplasmic reticulum that promotes disulfide bond formation in client proteins through its thiol-disulfide oxidase activity. Paip1 was more abundant in FLExDU4 as compared to WT mice, but FLExDUX4 did not change the abundance by VWR. Paip1 is known as a coactivator in the regulation of translation initiation of poly(A)-containing mRNAs.

**Figure 5.**
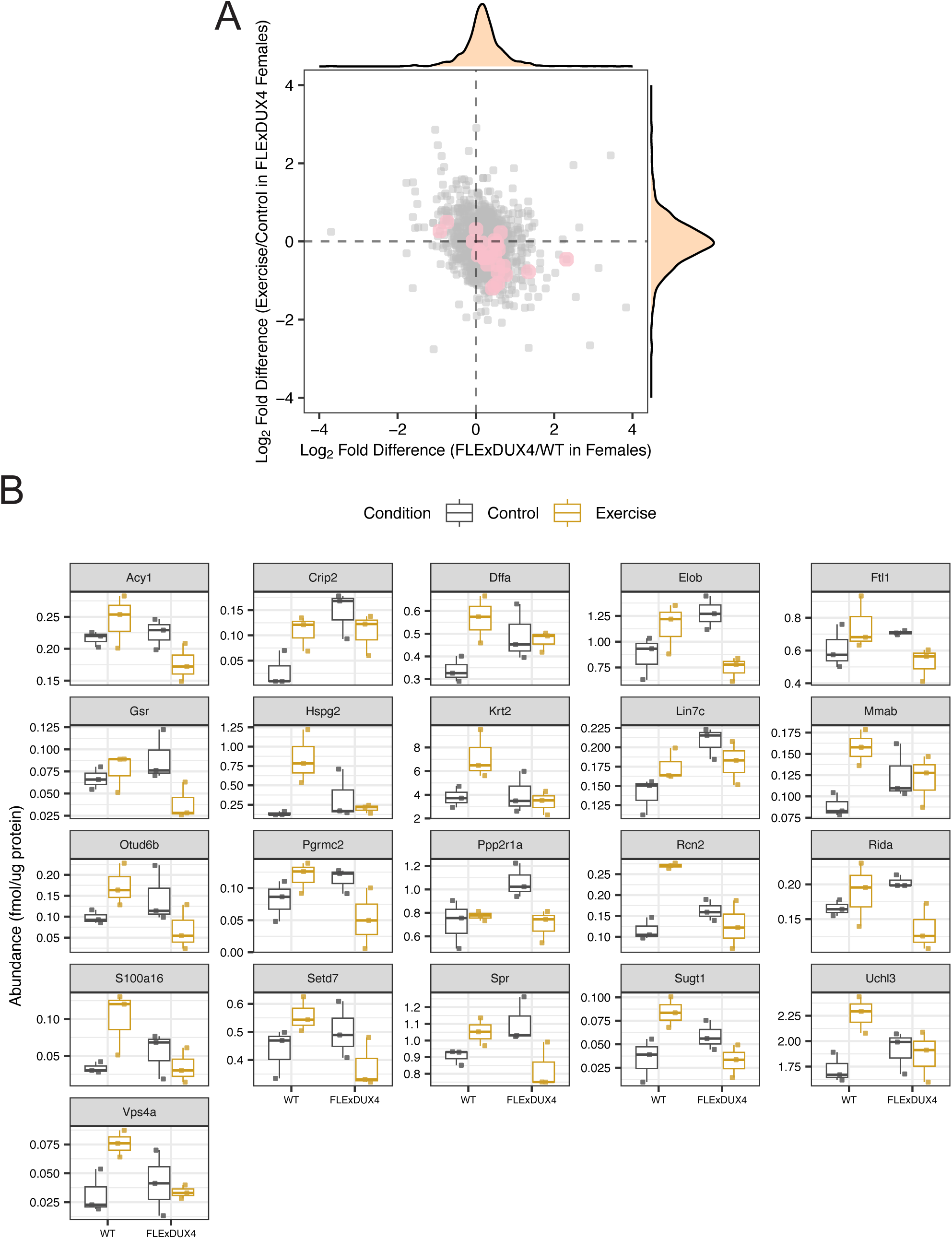
VWR normalized altered protein abundance involved in the ubiquitin proteasome system in female FLExDUX4 mice. *(**A**)* Scatter plot comparing the Log_2_ Fold-Difference between FLExDUX4 control and WT control (x-axis) and the Log_2_ Fold-Difference between FLExDUX4 Exercise and FLExDUX4 control (y-axis). Pink data points represent proteins with an interaction effect between training status and Genotype (P < 0.05) in female mice. Density plots display the distribution of individual protein abundance data both in X and Y axis. (***B***) Boxplot of 27 proteins that are localized in the bottom right quadrant with an interaction effect (P < 0.05, two-way ANOVA of training status and Genotype in female mice).

**Figure 6.**
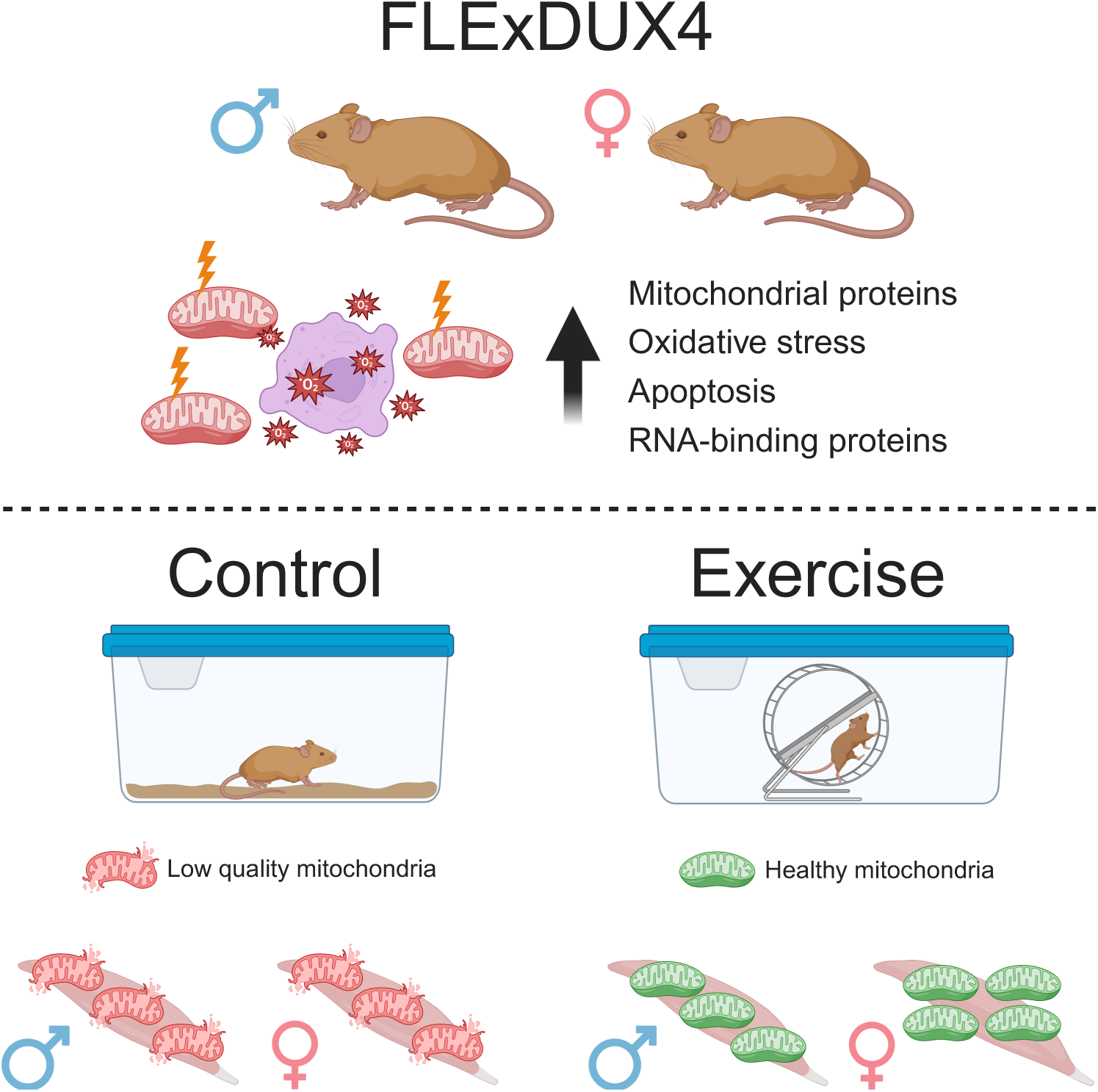
Muscle proteome in FLExDUX4 mice recapitulate key skeletal muscle clinical characteristics of FSHD with sexual dimorphism of mitochondrial proteome adaptation to VWR. Proteome in FLExDUX4 triceps brachii are characterized by altered proteome associated with mitochondria, oxidative stress, apoptosis. Moreover, RNA-binding proteins exhibit a sex-specific difference specifically in FLExDUX4 mice. These proteomic features of FLExDUX4 are consistent with our earlier proteomic study on myoblasts from people affected by FSHD (13) discovering an accumulation of potentially damaged or old mitochondrial proteins. In FLExDUX4 mice, the average mitochondrial proteins are specifically increased in females. In some individual mitochondrial proteins, sexual dimorphism of mitochondrial proteome adaptation to voluntary wheel running (VWR) was uncovered with males decreased, whereas females increased mitochondrial proteins by VWR. In contrast, this phenomenon was not observed in WT mice where both females and males commonly increased mitochondrial proteins by VWR. FLExDUX4 males respond positively to VWR, including increases in the expression of mitochondrial genes and myofibre oxygen consumption rates (26), indicating an improvement of mitochondrial quality by VWR in FLExDUX4 mice. Created with BioRender.com.

## Discussion

We investigated triceps brachii muscle of single transgene FLExDUX4 mice to generate new insights into the effects of chronic low-level DUX4 expression, which may reflect disease processes of FSHD in humans. To the best of our knowledge, this is the first report on the muscle proteome of FLExDUX4 mice. The muscle proteome of 6-month-old FLExDUX4 mice recapitulated key clinical characteristics of human FSHD including alterations to the mitochondrial proteome, RNA-binding proteins, apoptosis, oxidative stress, and proteostasis. The current data in FLExDUX4 mice align with our earlier proteomic study on myoblasts from people affected by FSHD (13) and add to our earlier transcriptomic analysis (26) on muscle responses to VWR in male FLExxDUX4 and WT mice. Our data point to atypical proteome adaptation to exercise specifically in male FLExDUX4 mice and support the validity of the FLExDUX4 murine model to recapitulate clinical features of chronic low levels of DUX4 expression characteristic of the human disease.

In both sexes, mitochondria proteins were more abundant in FLExDUX4 compared to WT, including mitochondrial ribosomal subunits (Mrpl12, Mrps28), subunits of respiratory chain Complex l (Ndufa8, Ndufb6), Complex ll (Sdhb), and Complex lll (Uqcrh), and mitochondrial import receptor subunit (Tomm6). In contrast, the mitochondrial proteome response to VWR was markedly different between female and male FLExDUX4 mice. In female FLExDUX4, the effect of VWR matched the pattern seen in WT mice and was associated with gains in the abundance of mitochondrial proteins, whereas in male FLExDUX4 VWR was associated with lesser abundance of mitochondrial proteins compared to non-exercised FLExDUX4 males. Bittel et al. (26) reports FLExDUX4 males do respond positively to VWR, including increases in the expression of mitochondrial genes and myofibre oxygen consumption rates in FLExDUX4 males but mtDNA abundance did not alter. Myoblasts from people with FSHD (13) exhibit a heightened abundance of mitochondrial proteins, including respiratory complexes and mitochondrial ribosome subunits that also had low rates of turnover, indicative of an accumulation of older or damaged proteins. Therefore, co-occurring gains in mitochondrial function alongside decreases in mitochondrial protein abundance in male FLExDUX4 may suggest exercise improves mitochondrial function by removing accumulated dysfunctional proteins.

FSHD-affected muscle exhibits mitochondrial dysfunction, including badly formed mitochondrial cristae and separation of the inner and outer membranes with evidence of protein aggregation by electron microscopy (10). Proteins involved in the mitochondrial contact site and cristae organising system (MICOS), including Chchd3 and Samm50 were more abundant in FLExDUX4 mice. Samm50 is a physical interactor of Chchd3 (43) and also interacts with ATG8 family proteins and p62/SQSTM1 to function as a receptor for mitophagy (44). Mitochondrial oxidative phosphorylation requires the interaction between p62/SQSTM1 and Samm50 to selectively deliver MICOS and SAM complex proteins to the lysosome to maintain mitochondrial proteostasis (44). Samm50 knock down cells exhibited abnormal and distorted mitochondrial cristae, suggesting a key function of Samm50 to maintain mitochondria (44). Optn is also a known autophagy receptor protein associated with the degradation of damaged mitochondria (45). The greater abundance of Chchd3, Samm50, Optn in FLExDUX4 may indicate a higher but unmet demand for mitophagy and suggest mitochondrial structural integrity or mitophagy flux might be impaired through the reduction of autophagosome formation, lysosomal fusion, or lysosomal degradation, which results in the accumulation of autophagy receptors due to incomplete clearance of damaged mitochondria. It is well established that muscle affected by DUX4 or FSHD has lower mitochondrial respiratory function (15, 26). As mitochondrial function is impacted by mitochondrial cristae (46), mechanisms that control mitochondrial cristae might be a potential therapeutic target in FSHD.

FSHD myoblasts are more susceptible to oxidative stress with a concomitant lower expression of some genes (e.g., GSTT2, GSR, HSPA4) involved in antioxidant processes (47). In our proteomic analysis, Gsr and Hspa4 were quantified, but did not differ between genotypes. In our proteomic analysis, Msrb2 was more abundant in FLExDXU4 mice. Msrb2 is a methionine sulfoxide reductase found in mitochondria, which catalyzes the reduction of free and protein-based methionine sulfoxides to methionine, and is involved in antioxidant mechanisms (48). Idh1 is essential to generate NADPH and an increase in Idh1 in FLExDUX4 may contribute to neutralizing reactive oxygen species from oxidative stress, as NADPH is known as a cofactor for the glutathione and thioredoxin systems. Peroxiredoxin 2 (Prdx2) a thiol-specific peroxidase to neutralize excessive ROS (49). Thioredoxin-like Protein 1 (Txnl1) has redox activities in disulfide reduction reactions, but Txnl1 was also found as an ATP-independent chaperone (50). Glutathione S-Transferase Omega-1 (Gsto1) is known to play a key role in glutathionylation (51). Increases in these proteins may implicate the higher demand of antioxidant defense mechanisms due to potentially increased oxidative stress in FLExDUX4 mice. These data also suggest a connection between mitochondrial dysfunction and elevated oxidative stress in FSHD.

Muscle affected by FSHD exhibit elevated apoptosis (52) and mitochondria contribute to the regulation of apoptosis via mitochondrial-derived cytochrome c (53). Fam162a was more abundant in FLExDUX4 mice and is a HIF-1α-responsive gene, which induces apoptosis via mitochondrial apoptotic cascades, such as cytochrome c release and caspase 9 activation (54). Indeed, Cycs, a well-known trigger of the intrinsic apoptotic pathway, was more abundant in FLExDUX4 mice. Furthermore, Dffa was increased in FLExDUX4, and this protein is known a substrate of Caspase-3 (55). Hint 2 was more abundant in FLExDUX4 mice and overexpression of Hint2 was shown to increase apoptosis (56). Eif5a is a translation factor required for translation elongation and termination. In our proteomic data, Eif5a was more abundant in FLExDUX4 and overexpression of Eif5a increased ROS and induced apoptosis (57). Additionally, Eif5a is also known to translate ATG3 protein (58) that facilitates LC3B lipidation and autophagosome formation.

DUX4 activation disrupts RNA metabolism, such as RNA splicing, surveillance, and transport pathways (59). Campbell et al. (20) showed that DUX4 induction disturbed nonsense-mediated RNA decay, which led to the accumulation of truncated RNA binding proteins and cytotoxicity associated with the truncated splicing factor SRSF3. In our proteomic data, two RNA binding proteins, Rbm3 and Rbm8a exhibited sex-specific differences in FLExDUX4 mice. Dresios et al. (60) reports Rbm3 directly interacts with 60S ribosomal subunits in an RNA-independent manner in rat brain and protein synthesis was increased 3-fold in cells expressing the Rbm3. Neither the size nor distribution of the polysome peaks was changed, instead microRNA in cells expressing the Rbm3 fusion protein was less abundant, and Rbm3 may modulate protein synthesis via a posttranslational mechanism in cooperation with microRNAs (60). We quantified serine/arginine-rich splicing factors (Srsf1, Srsf2, Srsf3, Srsf4, Srsf7) and Srsf4, which plays a role in alternative splice site selection during pre-mRNA splicing, was significantly more abundant in FLExDUX4 as compared to WT. Mitochondrial transcription rescue factor 1 (Mtres1) was more abundant in FLExDUX4 and an increase in Mtres1 was previously shown to have the RNA binding activity and prevent mitochondrial transcript loss under stress conditions (61). Accumulation of RNA-binding proteins in FLExDUX4 mice and a sex-specific differences in Rbm3 and Rbm8a might be key contributing factors to the disease mechanisms in FSHD.

Our work builds molecular evidence for the use of exercise as both a therapy in FSHD patients and as an experimental tool for discovering new pharmacological targets via animal models such as FLExDUX4. In people with FSHD, greater levels of physical activity in youth correlate with less severe clinical presentation in older age (63) and aerobic exercise training can bring significant gains in whole body oxygen uptake (64). Nevertheless, adherence to regular physical activity is low in people with FSHD (65) and a Cochrane Review of 14 trials (428 participants) concluded only that, despite no evidence of benefit, moderate intensity strength training in people with FSHD does not cause harm (66). Notwithstanding the lack of clear evidence on gross physiological benefits of exercise further, more detailed, molecular analysis on exercise responses in FSHD and DUX4 expressing muscle is warranted. In particular, our work supports the hypothesis that exercise benefits DUX4 expressing muscle by overcoming inadequacies in degradative processes that allow damaged mitochondria to accumulate.

## Conclusions

FLExDUX4 single transgenic mouse exhibits molecular features of FSHD pathology that align with proteomic findings in myoblasts from people affected by FSHD. Sexual dimorphism exists in FLExDUX4 muscle and particularly in the response of FLExDUX4 muscle to exercise, including core disease processes of mitochondrial dysfunction and disruption to RNA processing. Exercise reversed proteome features of FLExDUX4 mice in both sexes despite the fact that the effect of FLExDUX4 differed between sexes. Our data highlight the importance of identifying sex-specific diagnostic biomarkers for more reliable development of FSHD biomarkers and therapeutic targets. Our data provide a new hypothesis-generating resource for future mechanistic studies using a FLExDUX4 murine model of FSHD, including studies on mitochondrial proteostasis and muscle aerobic capacity in FSHD pathology.

## Abbreviations

Ambic: ammonium hydrogen bicarbonate
BSA: bovine serum albumin
DTT: dithiothreitol
DUX4: double homeobox 4
DUX4-fl: full-length DUX4 transgene
FASP: Filter-Aided Sample Preparation
FSHD: facioscapulohumeral muscular dystrophy
GO: gene ontology
HSP: heat shock proteins
mtDNA: mitochondrial DNA
PCA: principal component analysis
ROS: reactive oxygen species
TFA: trifluoracetic acid
VWR: voluntary wheel running

## Data availability

The mass spectrometry proteomics data generated in this study have been deposited to the ProteomeXchange Consortium via the PRIDE (67) with the dataset identifier PXD061097 and 10.6019/PXD061097.

## Supplemental data

This article contains supplemental data.

