## supplemental data for "Proteomic profiling uncovers sexual dimorphism in the muscle response to wheel running exercise in the FLExDUX4 murine model of facioscapulohumeral muscular dystrophy"

**Running title:** Proteomic analysis of triceps brachii muscle in FLExDUX4 mice

**Keywords:**

FSHD; DUX4; skeletal muscle; proteomics; exercise; mitochondria; RNA-binding proteins; apoptosis; oxidative stress

### Supplemental Figures

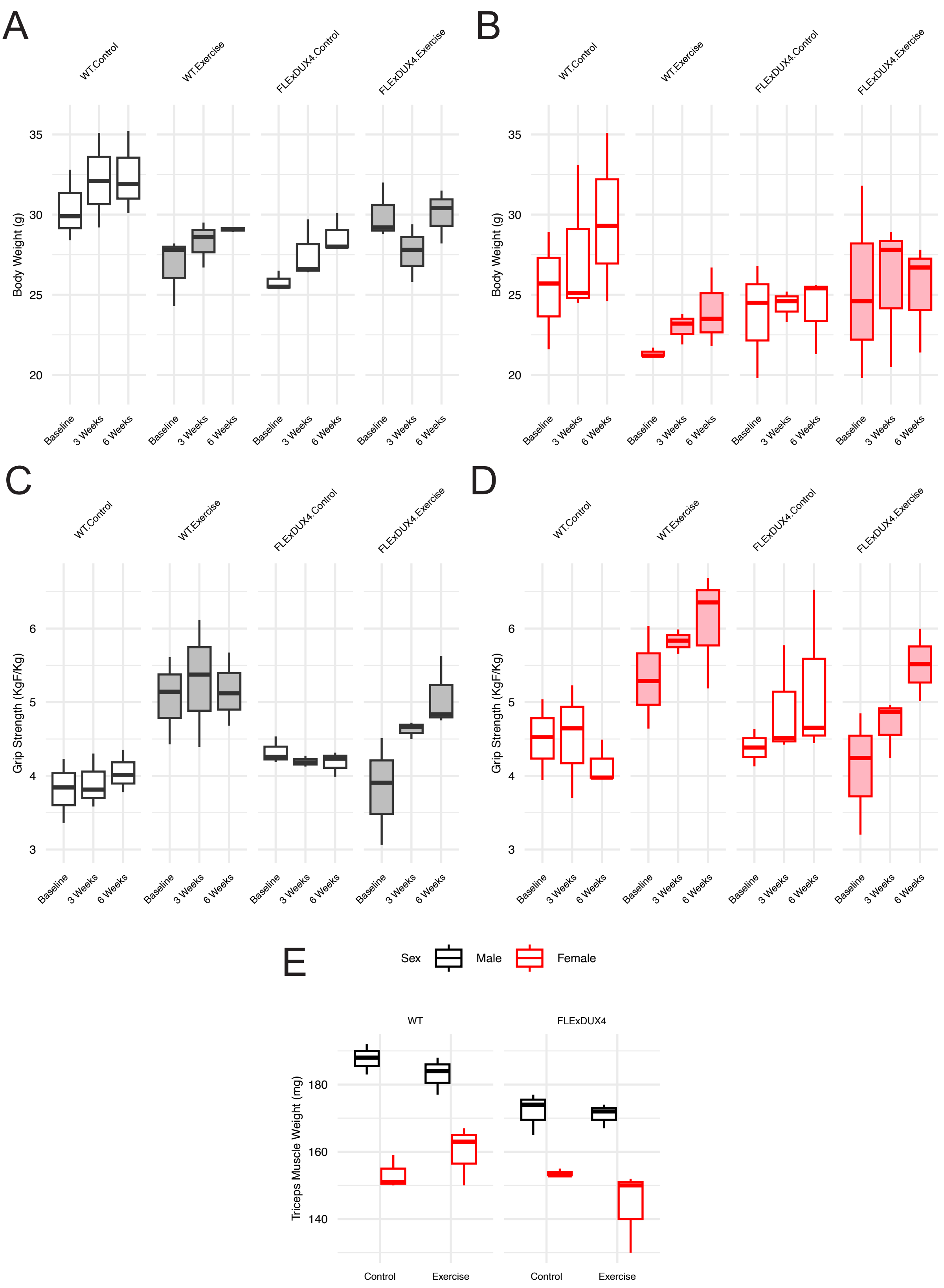


**Figure S1: FLExDUX4 mice increased grip strength in response to VWR without changing body weight and triceps brachii muscle mass**

Changes in body weight (g) among WT and FLExDUX4 male (**A**) and female (**B**) mice in the control and exercise groups. Changes in body-weight-normalized grip strength (KgF/Kg) among WT and FLExDUX4 male (**C**) and (**D**) female mice in the control and exercise groups. (**E**) Muscle weight of the triceps brachii (mg) at a 6-week time point.

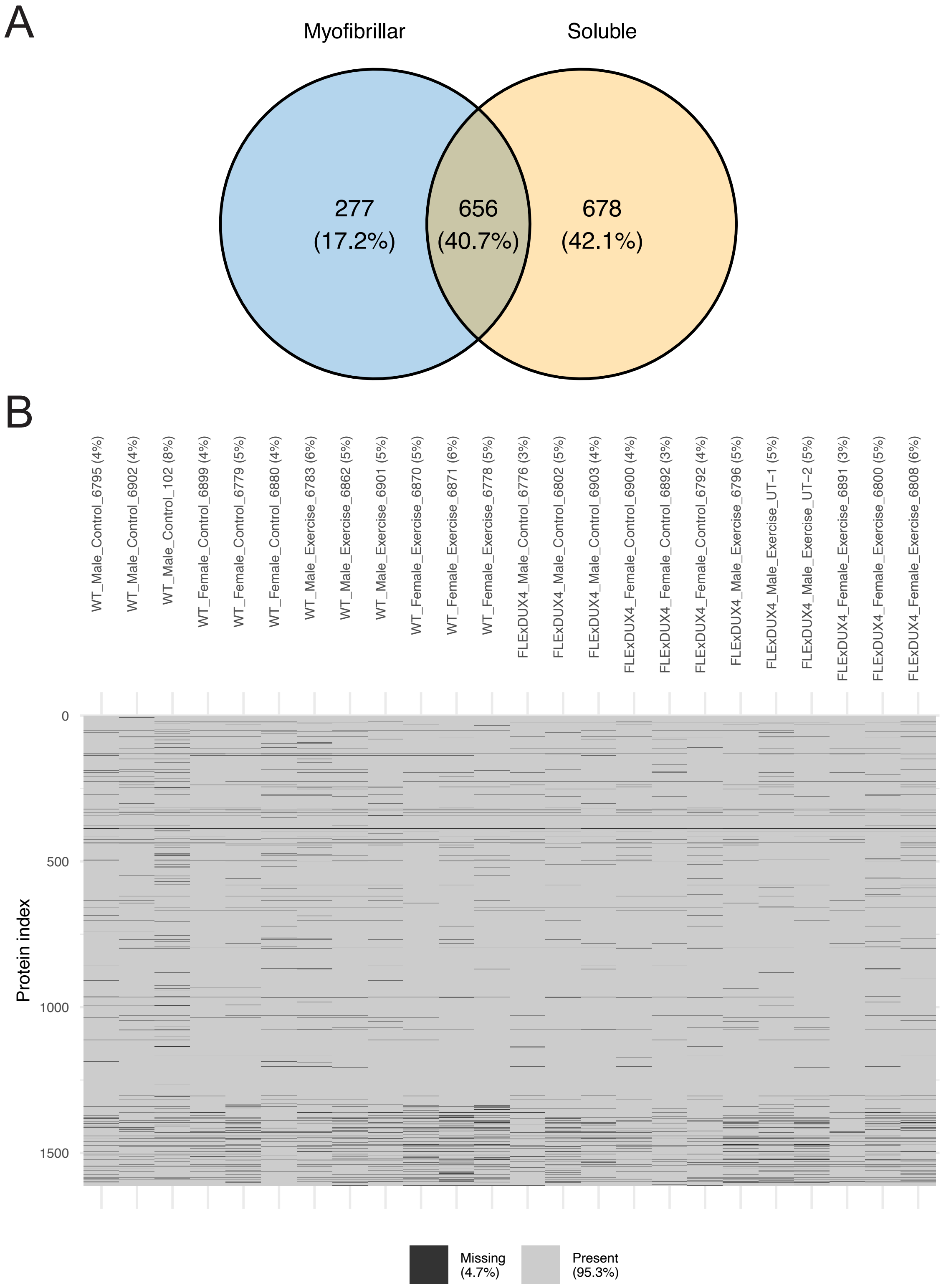


**Figure S2: Overlap of proteins between soluble and myofibrillar fractions and missing value distribution per sample.** (**A**) A Venn diagram comparing the number of proteins identified between soluble and myofibrillar fractions prior to missing values filtering. (**B**) A matrix heatmap visualizing missing values after summing data from soluble and myofibrillar fractions. Each column represents a sample group, and each row represents protein accession (n = 1611 proteins).

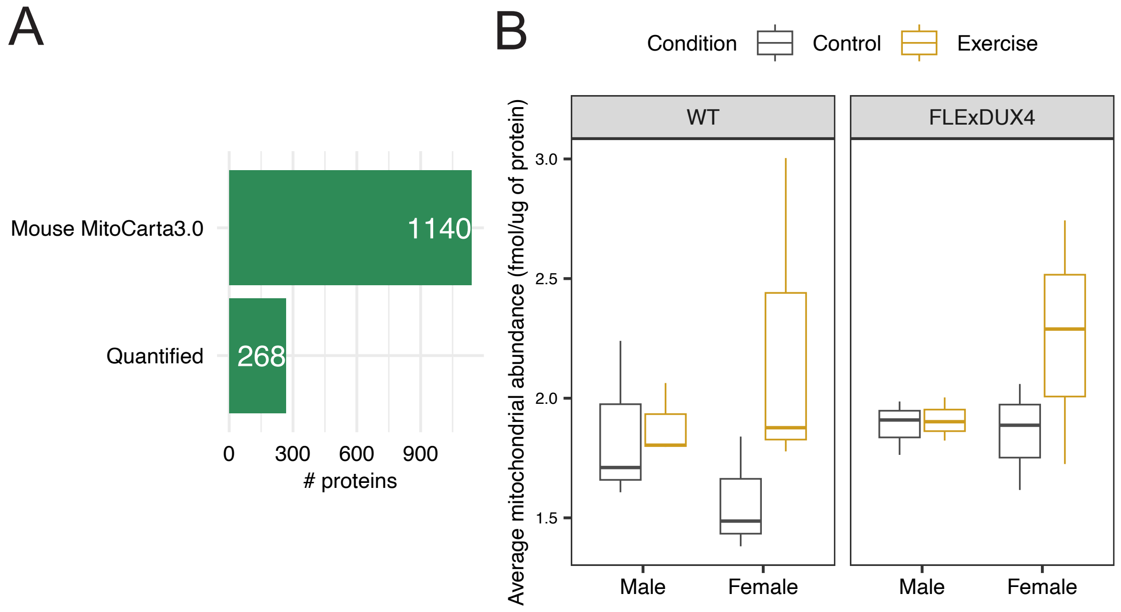


**Figure S3: Average mitochondrial protein abundance**

(**A**) Bar chart showing the number of proteins identified in Mouse MitoCarta3.0 and quantified in the current proteomic analysis. (**B**) Box plots comparing the average mitochondrial protein abundance (268 proteins) between genotype (WT vs FlexDUX4), sex (Male vs Female), and condition (Control vs Exercise). Three-way ANOVA did not exhibit statistical differences (P>0.05).

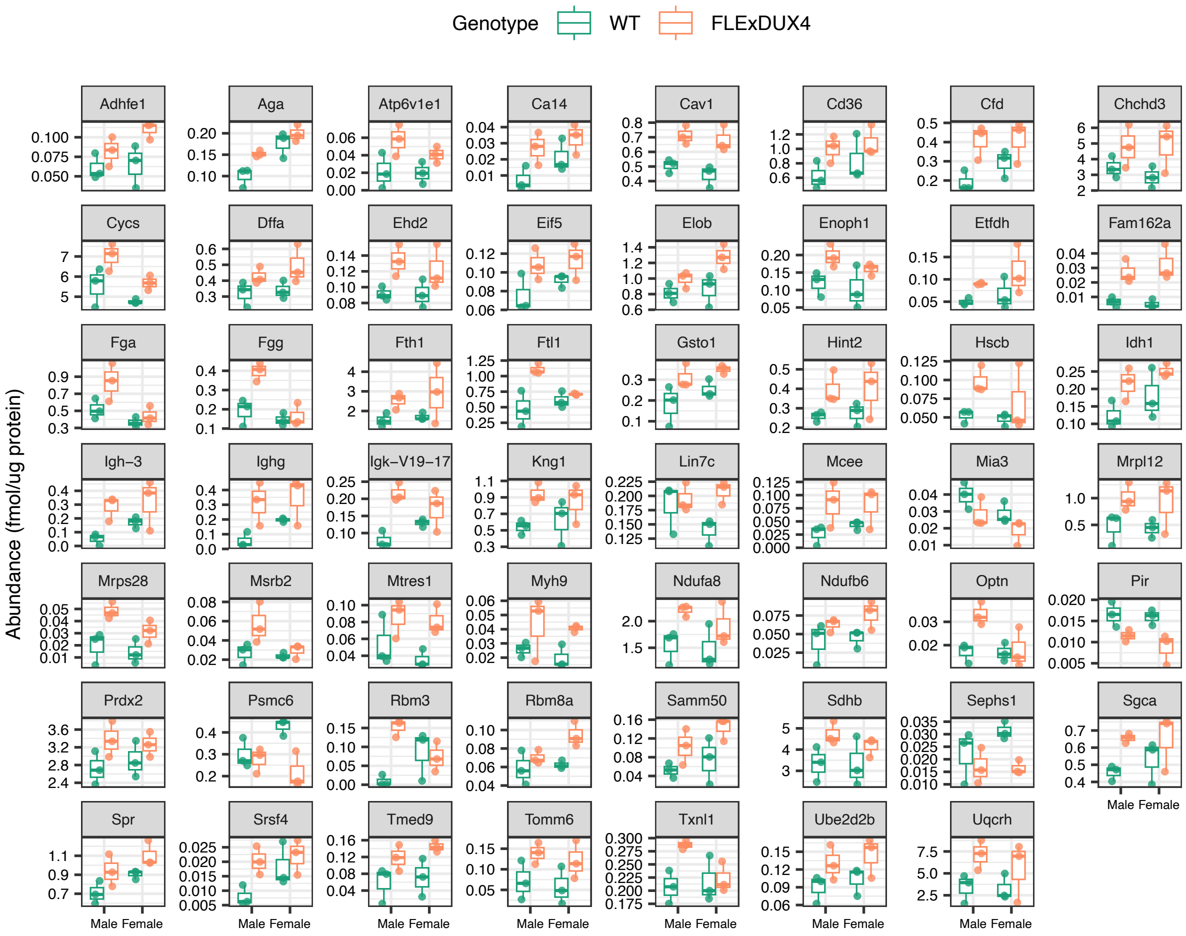


**Figure S4: Box plots of 55 proteins with a main effect of Genotype – Related to Figure 2**

Box plots illustrating differences of protein abundance between Sex (Male versus Female) and Genotype (FLExDUX4 versus WT) in FLExDUX4 mice.

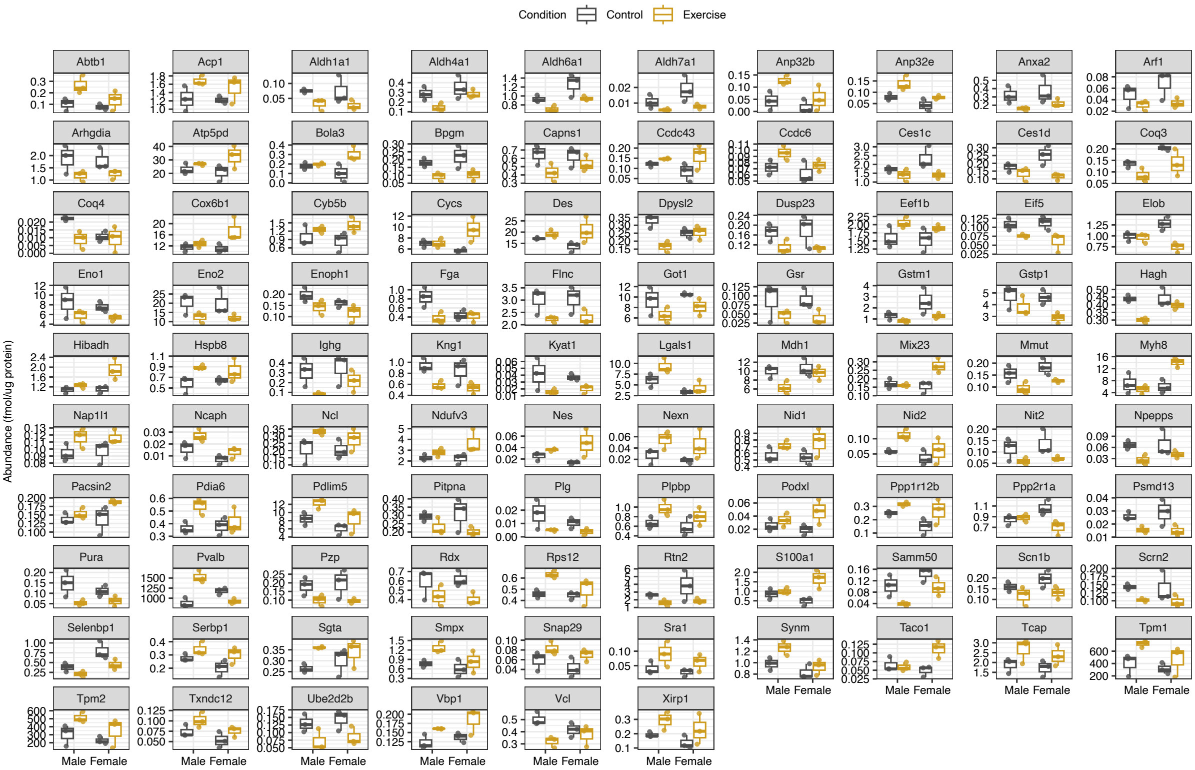


**Figure S5: Box plots of 96 proteins with a main effect of Condition – Related to Figure 3**

Box plots illustrating differences of protein abundance between Sex (Male versus Female) and Condition (Exercise versus Control) in FLExDUX4 mice.

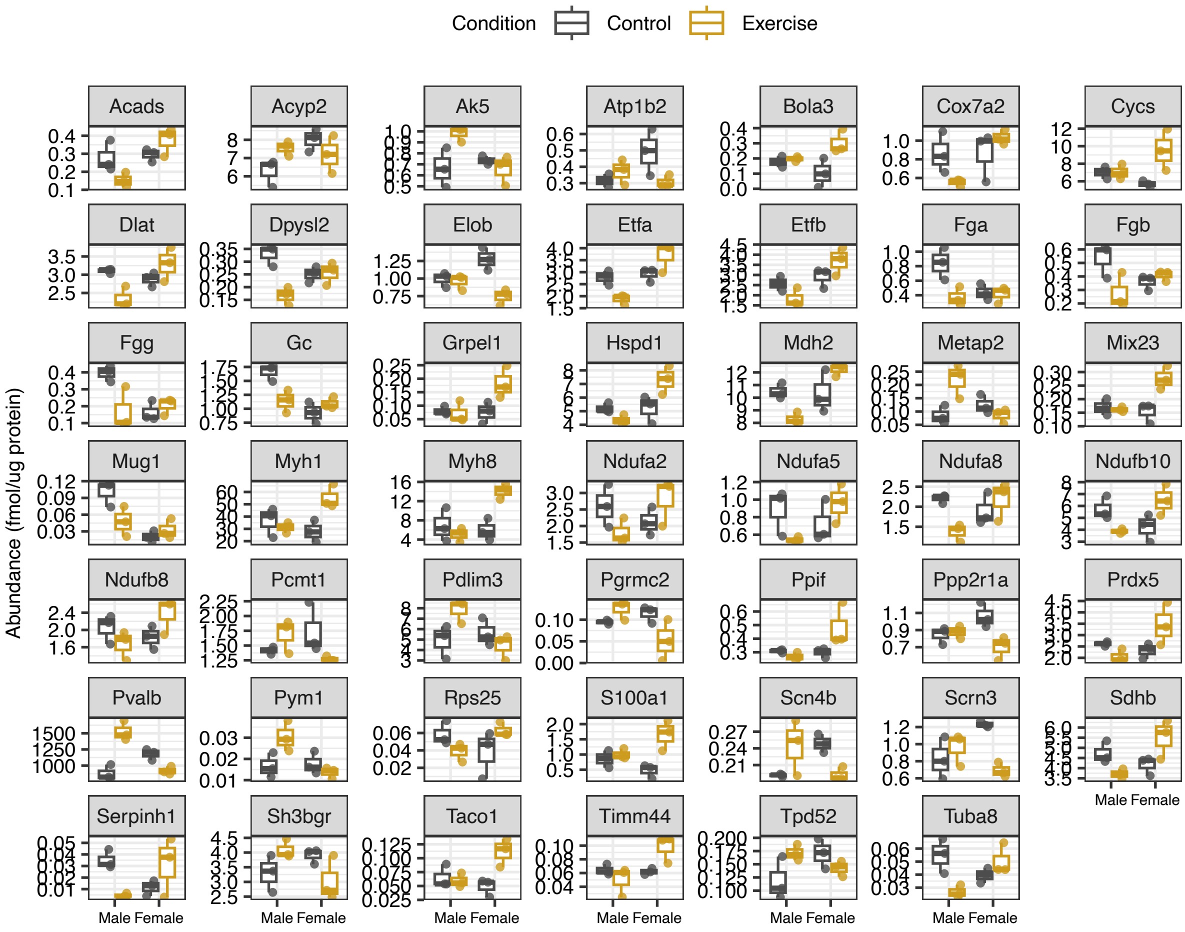


**Figure S6: Box plots of 96 proteins with an interaction effect – Related to Figure 3**

Box plots illustrating differences of protein abundance between Sex (Male versus Female) and Condition (Exercise versus Control) in FLExDUX4 mice.

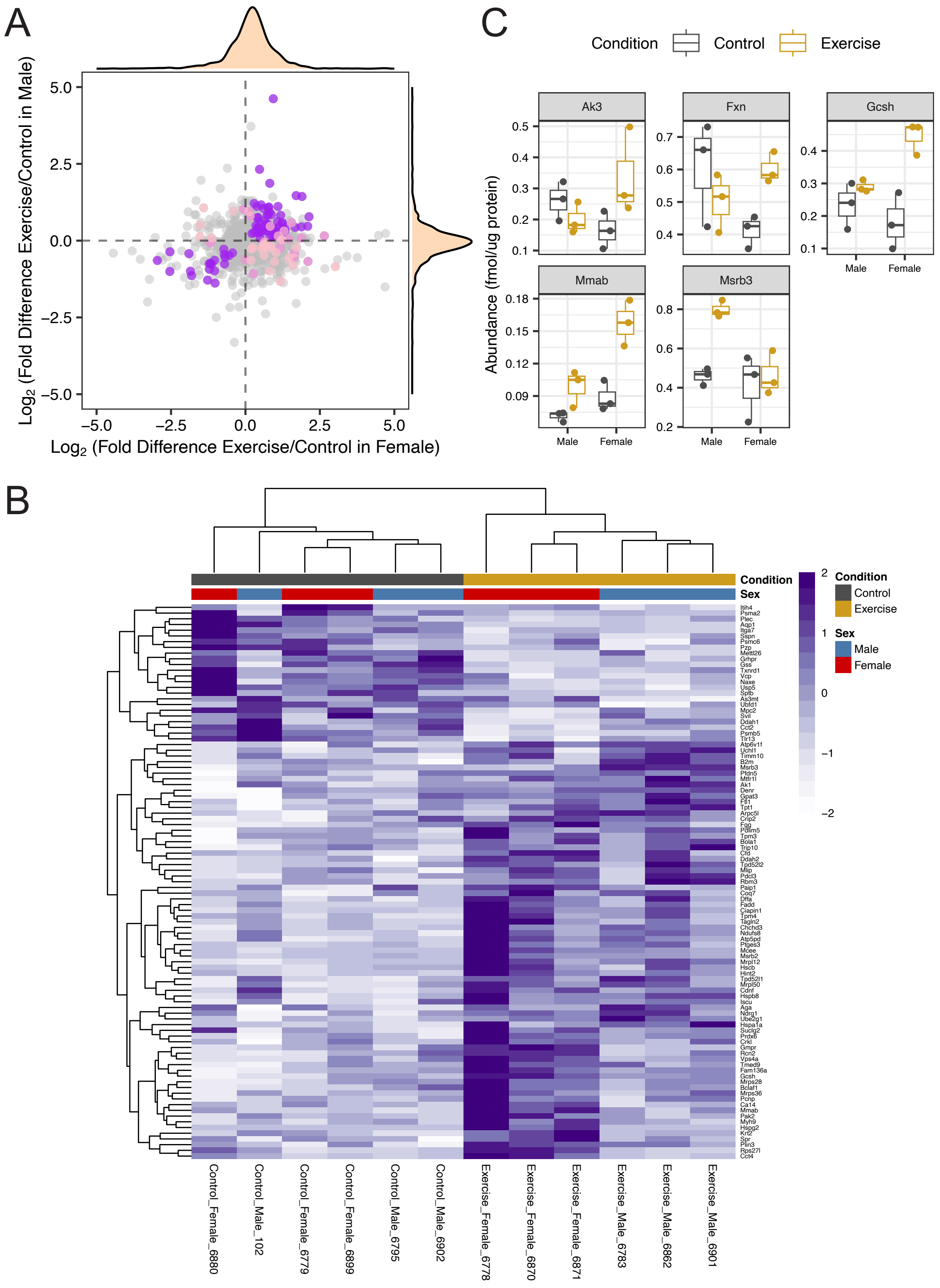


**Figure S7: WT mice does not exhibit sexual dimorphism of mitochondrial proteome adaptation to voluntary wheel running**

(**A**) Scatter plot comparing the differences in the $Log_2$ Fold-Difference (Exercise/Control) between protein abundance of females (x-axis) and males (y-axis) in WT mice. Purple data points represent proteins with a main effect of Genotype ($P < 0.05$) and pink data points represent proteins with an interaction effect between Genotype and Sex ($P < 0.05$). Density plots display the distribution of individual protein abundance data both in X and Y axis. (**B**) Heatmap of proteins with significant differences in abundance between Exercise and Control muscle (97 proteins, a main effect of Condition; $P < 0.05$). Protein abundances were normalized by row and represented by colors according to the scale in the key. Both rows (proteins) and columns (samples) were clustered using hierarchical clustering. (C) Boxplot of 5 proteins filtered by Mouse MitoCarta3.0 that exhibit an interaction effect ($P < 0.05$, two-way ANOVA of Sex and Condition in WT mice).

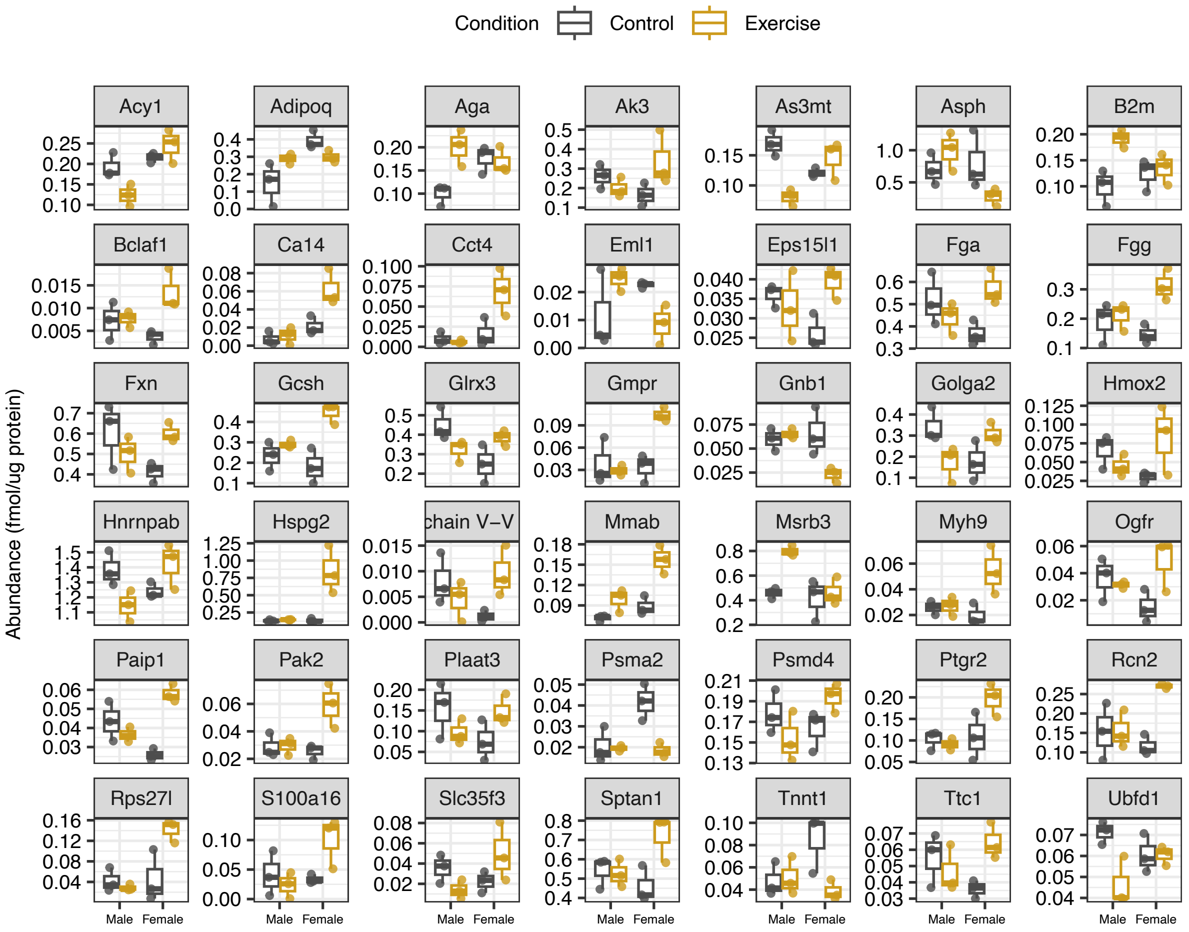


**Figure S8: Box plots of 42 proteins with an interaction effect – Related to Figure S7**

Box plots illustrating differences of protein abundance between Sex (Male versus Female) and Condition (Exercise versus Control) in WT mice.

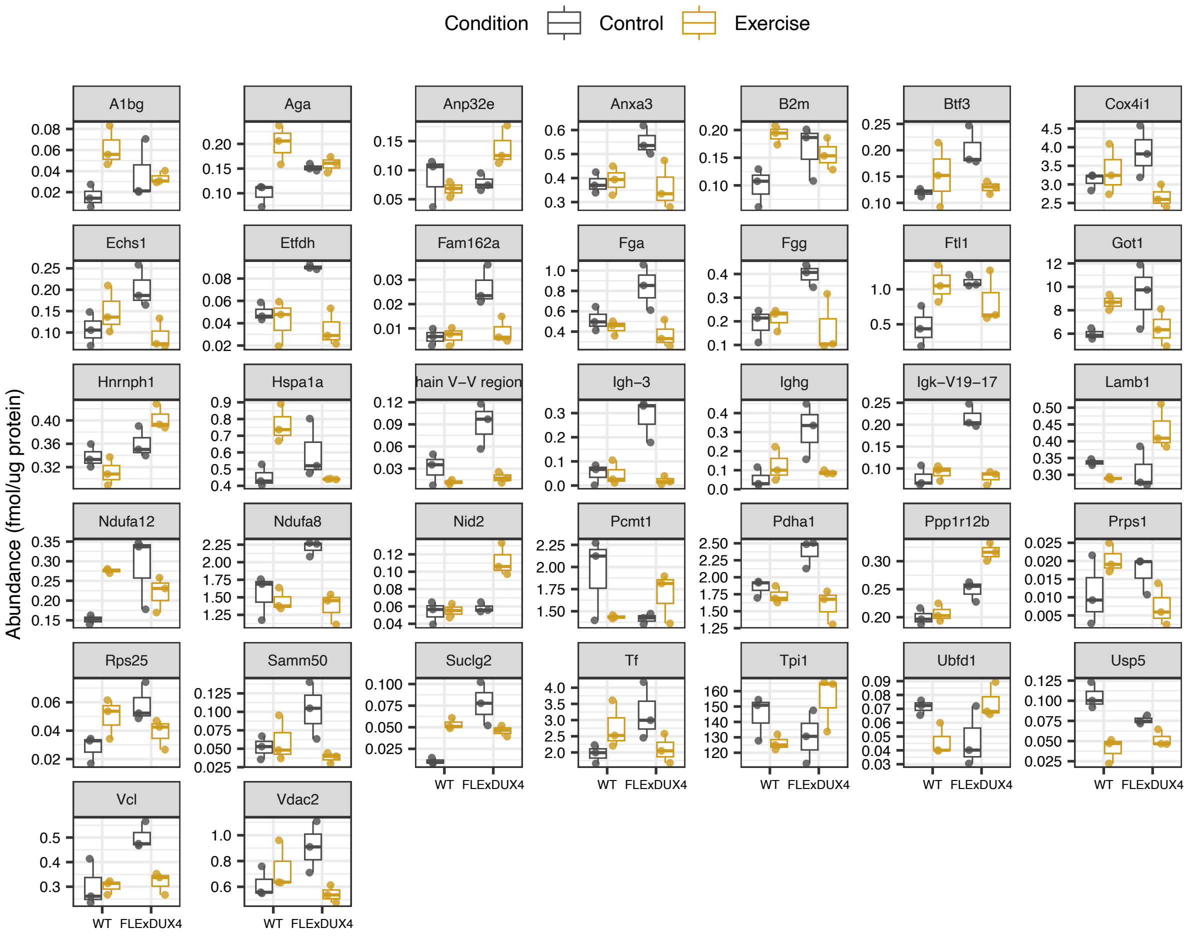


**Figure S9: Box plots of 37 proteins with an interaction effect – Related to Figure 4**

Box plots illustrating differences of protein abundance between Genotype (FLExDUX4 versus WT) and Condition (Exercise versus Control) in male mice.

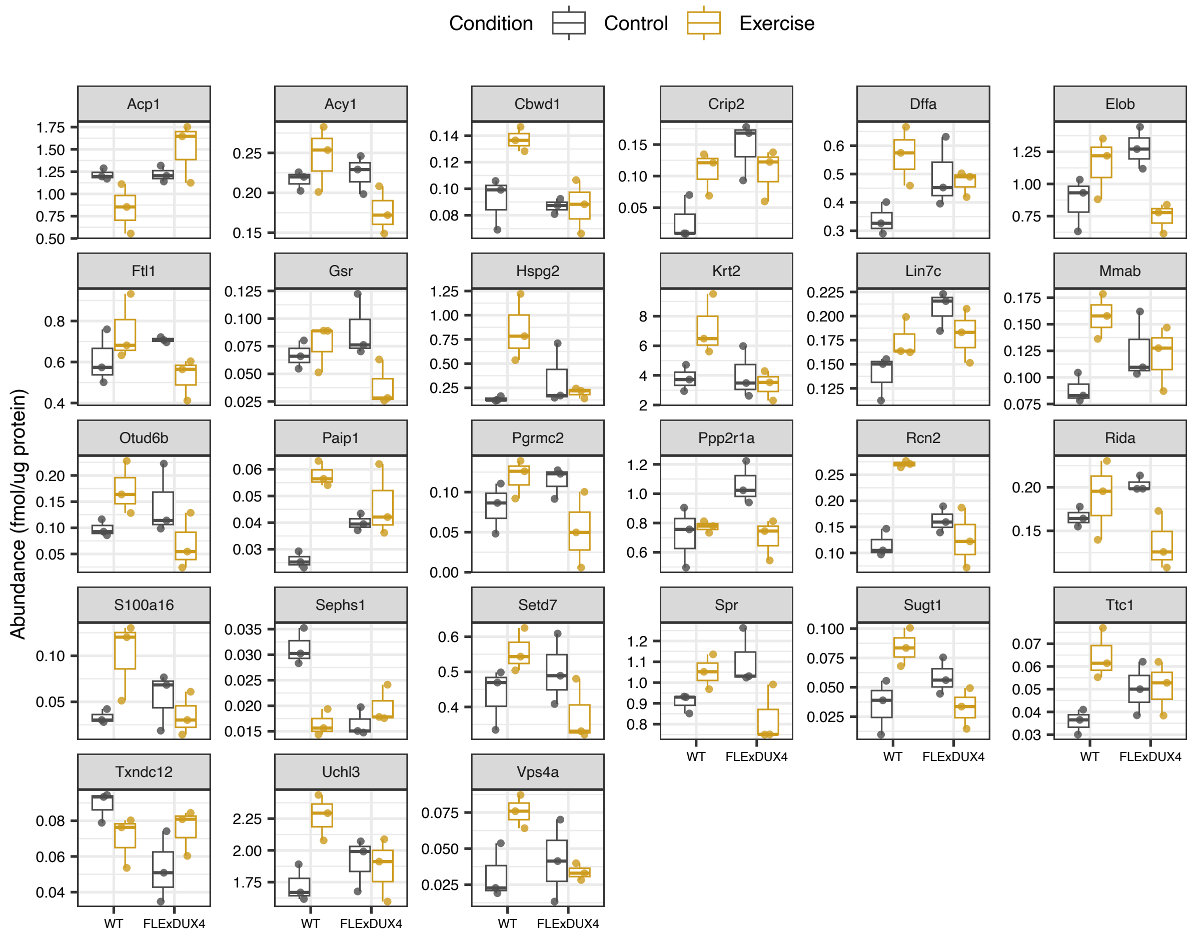


**Figure S10: Box plots of 27 proteins with an interaction effect – Related to Figure 5**

Box plots illustrating differences of protein abundance between Genotype (FLExDUX4 versus WT) and Condition (Exercise versus Control) in female mice.
